# Mitochondrial dysfunction and oxidative stress contribute to cognitive and motor impairment in FOXP1 syndrome

**DOI:** 10.1101/2021.07.05.451143

**Authors:** Jing Wang, Henning Fröhlich, Felipe Bodaleo Torres, Rangel Leal Silva, Amit Agarwal, Gudrun A. Rappold

## Abstract

There is increasing evidence that mitochondrial homeostasis - influenced by both genetic and environmental factors - is crucial in neurodevelopment. FOXP1 syndrome is a neurodevelopmental disorder that manifests motor dysfunction, intellectual disability, autism and language impairment. In this study, we used a *Foxp1*^+/−^ mouse model to address whether cognitive and motor deficits in FOXP1 syndrome are associated with mitochondrial dysfunction and oxidative stress. Here we show that genes with a role in mitochondrial biogenesis and dynamics (e.g. *Foxo1*, *Pgc-1α*, *Tfam*, *Opa1*, and *Drp1)* were dysregulated in the striatum of *Foxp1*^+/−^ mice at different postnatal stages. Furthermore, in the striatum of *Foxp1*^+/−^ animals, mitochondrial membrane potential was disrupted, and reactive oxygen species, lipid peroxidation and cytochrome c release were significantly elevated. These features can explain the reduced neurite branching, learning and memory, endurance, and motor coordination that we observed in these animals. Taken together, we provide strong evidence of mitochondrial dysfunction in *Foxp1*^+/−^ mice, suggesting that insufficient energy supply and excessive oxidative stress underlies the cognitive and motor impairment in FOXP1 deficiency.

## Introduction

Autism spectrum disorder (ASD) is a neurodevelopmental disorder characterized by impaired social communication, restricted or stereotypical interests, and repetitive behaviors. Molecular genetic studies on ASD individuals and their families have identified several hundred ASD risk genes, including the forkhead box protein P1 (*FOXP1*) gene. In humans, haploinsufficiency of *FOXP1* leads to FOXP1 syndrome (OMIM 605515), a neurodevelopmental disorder characterized by delayed motor and language development, intellectual disability, autistic traits and dysmorphic features [1, 2]. Since the first report in 2010 [1], more than hundred cases with *de novo* mutations have been described in the literature, and *FOXP1* is now among the highest ranked ASD genes according to the SFARI Gene database [3].

FOXP1 is a member of the Forkhead Box P (FOXP) subfamily of transcription factors, which also includes FOXP2, FOXP3, and FOXP4. Both FOXP1 and FOXP2 are highly conserved in vertebrates and form homo- and heterodimers [4, 5], and while *FOXP2* mutations have been linked with a distinct speech and language disorder [6], *FOXP1* mutations affect much more global neural functions. They may also form oligomers with FOXP4 [5], suggesting that these three related transcription factors regulate cognitive behavior and motor development via common pathways [7].

The complete loss of *Foxp1* in mice is lethal at E14.5 [8], but the phenotype of heterozygous *Foxp1*^+/−^ mice resembles that of individuals with FOXP1 syndrome. *Foxp1*^+/−^ animals exhibit behavioral and muscle deficits such as reduced neonatal ultrasonic vocalization and grip strength [9], that are first indications for social deficits and hypotonia in animals. Furthermore, *Foxp1*^+/−^ mice display esophageal achalasia and impaired peristalsis in the colon [10], reflecting the gastrointestinal disturbances seen in many individuals with FOXP1 syndrome [2, 11].

In the brain, Foxp1 is mainly expressed in the cortex, hippocampus and striatum [7, 12]. In the cortex, Foxp1 is mainly expressed in the intratelecephalic projection neurons in layers 3-5, regulating neonatal vocalizations [13]. In the hippocampus, Foxp1 expression is seen in the CA1 and subiculum region, required for spatial learning and synaptic plasticity [14]. In the striatum, Foxp1 has been shown its most abundant expression and to regulate the cellular composition and connectivity in a cell type- specific manner [15]. Thus, Foxp1 has an essential role in regulating neurogenesis and synapse organization. However, the role of Foxp1 and its transcriptional regulators have not been studied on metabolic disturbances and mitochondrial malfunction. In fact, mitochondrial function plays an important role in neurogenesis, synaptic plasticity, and neurotransmission [16, 17, 18].

Metabolic disturbances and mitochondrial malfunctions are also more common in individuals with ASD than in the general population, indicating that they play a role in the pathogenesis of ASD [19, 20]. However, only certain subgroups of the ASD population are affected. To investigate whether mitochondrial impairment might underlie the cognitive and motor features of FOXP1 syndrome, we analyzed mitochondrial structure and motility, and the expression of genes that regulate mitochondrial biogenesis and dynamics in the striatum of *Foxp1*^+/−^ mice. We examined autophagy, production of reactive oxidative species (ROS), metabolic status and cytochrome c release in these animals. In addition, we analyzed neuronal morphology, motor function, as well as emotional and spatial learning and memory.

## Methods and Materials

### Animals

*Foxp1*^+/−^ mice [8] were backcrossed with C57BL/6J mice for at least 12 generations to obtain congenic animals. Mice were kept in a specific pathogen-free Biomedical Animal Facility under a 12 h light-dark cycle and given *ad libitum* access to water and food. All procedures were conducted in compliance with the National Institutes of Health Guidelines for the Care and Use of Laboratory Animals and were in accordance with the German Animal Protection Law (TierSCHG). The day of birth was considered as postnatal day (P) 0.5. Animal studies were approved by the Governmental Council Karlsruhe, Germany (license number: 35.9185.81-G-100/16 and 35.9185.81-G-102/16).

### Behavioral analysis

CatWalk gait analysis, treadmill exhaustion test and active place avoidance were performed as described in the Supplementary information.

**Analysis of mRNA and protein expression levels as well as the calculation of mitochondrial DNA copy number and DNA deletion are** described in the Supplementary information. Primer sequences are listed in Supplementary Table 1, primary and secondary antibodies in Supplementary Table 2.

### Detection of mitochondrial membrane potential

Mitochondrial membrane potential of primary neurons was measured by fluorescent staining of Tetramethylrhodamine methyl ester (Image-iT™ TMRM Reagent, ThermoFisher Scientific, Waltham, MA, USA) according to the manufacturer’s instructions.

### Measuring reactive oxygen species (ROS) in striatal tissue

Striatal ROS was marked by 2’ 7’-dichlorodihydrofluorescein diacetate (Molecular Probes™ H2DCFDA (H2-DCF, DCF), ThermoFisher Scientific) as previously described [21] and assessed by DS-11 Series Spectrophotometer/Fluorometer (DeNovix Inc., Wilmington, DE, USA).

### Detection of lipid peroxidation products

Lipid peroxidation products were analyzed in striatal tissue using the Bioxytech LPO-586 kit (OxisResearch™, Burlingame, CA, USA) according to the manufacturer’s instructions. The tissue was homogenized in PBS-containing butylated hydroxytoluene (BHT) to prevent sample oxidation during homogenization.

### Isolation of mitochondria and cytochrome c release

Mitochondria from striatal tissue were isolated as previously reported [22], except minor changes (Supplementary information). Cytochrome c release was determined by calculating the ratio of occurrence in the mitochondria to cytoplasm. The isolation of mitochondrial and cytosolic proteins was conducted as previously described [22]. Cytochrome c, Cox IV (a marker for mitochondrial proteins) and Gapdh (a marker for cytosolic proteins) were evaluated by western blot.

### Staining of mitochondrial superoxide in living cells

Superoxides were visualized by MitoSOX™ Red reagent (ThermoFisher Scientific) according to the manufacturer’s instructions. In brief, primary striatal neurons were stained with 100 nM MitoSOX™ Red reagent for 30 min at 37 °C and examined by the Automated Inverted Microscope DMI4000B (Leica Camera AG, Wetzlar, Germany).

### Mitochondrial dynamics and structural analyses

Striatal neurons were incubated with MitoTracker Red CMXRos (Cat M7512. ThermoFisher) at DIV8. Live-cell imaging was performed using a custom built two-photon microscope (Bergamo II, Thorlabs, Newton, New Jersey, USA) using a Nikon NIR, 60X 1.0 NA objective (details in Supplementary information). For the mitochondrial dynamics analyses, kymographs were generated on the time series images using the Icy Kymograph Tracker tool (version 2.1.2.0) [23, 24]. For the mitochondria structure analysis, single frame images were segmented with Ilastik (version 1.4.0b7) [25] and surface area was measured with the *Region Properties* MATLAB function (Mathworks, Natick, Massachusetts, USA, 2020a).

**Primary striatal neuron culture, Sholl analysis, co-immunoprecipitation (Co-IP), cellular adenosine compound measurements and Oxygen Consumption Rate (OCR) and Extracellular Acidification Rate (ECAR)** were performed using standard protocols (Supplementary information).

### Statistics

Mice were randomly assigned to tests and investigators were blinded to genotypes. IBM SPSS Statistics 25 and Microsoft Office Excel software were used to analyze primary data. Outliers were determined via IBM SPSS Statistics 25 and strong outliers (≥3 standard deviations above mean) were excluded from further analysis. All data were checked for normal distribution via the Kolmogorov-Smirnov and Shapiro-Wilk test. Two-way ANOVA was performed with the litter as a second factor. In the analysis of mitochondrial dynamics, Mann-Whitney test was applied. *p*-values of ≤ 0.05 were considered significant.

## Results

### Dysregulated mitochondrial biogenesis in the *Foxp1*^*+/−*^ striatum decreases mitochondrial copy numbers and increases mitochondrial DNA deletions

Nervous system-specific and conventional *Foxp1* knockout mice [9, 12] and MRI images of FOXP1 syndrome individuals [2, 26] have revealed that the striatum is particularly affected by FOXP1 deficiency. To evaluate Foxp1 expression at early postnatal development as well as adult stage in the striatum, we compared *Foxp1* mRNA and protein level of *Foxp1*^+/−^ to that of WT littermates at postnatal day (P) 1.5, P12.5, and 8 weeks. The rational for the choice P1.5 is that at this time developmental stage the excitatory inputs to the striatum is minimal, by P12.5 the excitatory inputs to the striatum peaks and by 8 weeks striatal circuits are fully mature and the mice have complete motor function [27, 28]. Additionally, this maturation is accompanied by a significant decrease in *Foxp1* mRNA and protein with a 37% reduction of mRNA levels and 56% reduction of protein levels (both Foxp1 A and Foxp1 D) in adult *Foxp1*^+/−^ mice compared to WT animals (Fig. 1a and Supplementary Fig. 1a, 2a). *Foxo1*, a target of Foxp1 in the striatum [10, 12], was also significantly lower on mRNA and protein level at all three time points analyzed (Fig. 1b and Supplementary Fig. 1a, 2b). Foxp1 and Foxo1 interact in liver tissue and bind competitively to the insulin response element[29], our immunoprecipitation experiments did not reveal a physical interaction between Foxp1 and Foxo1 in striatal tissue, thereby suggesting that a direct Foxp1-Foxo1 interaction does not contribute to the striatal dysfunction in Foxp1-deficient mice (Supplementary Fig. 1b).

**Fig. 1:**
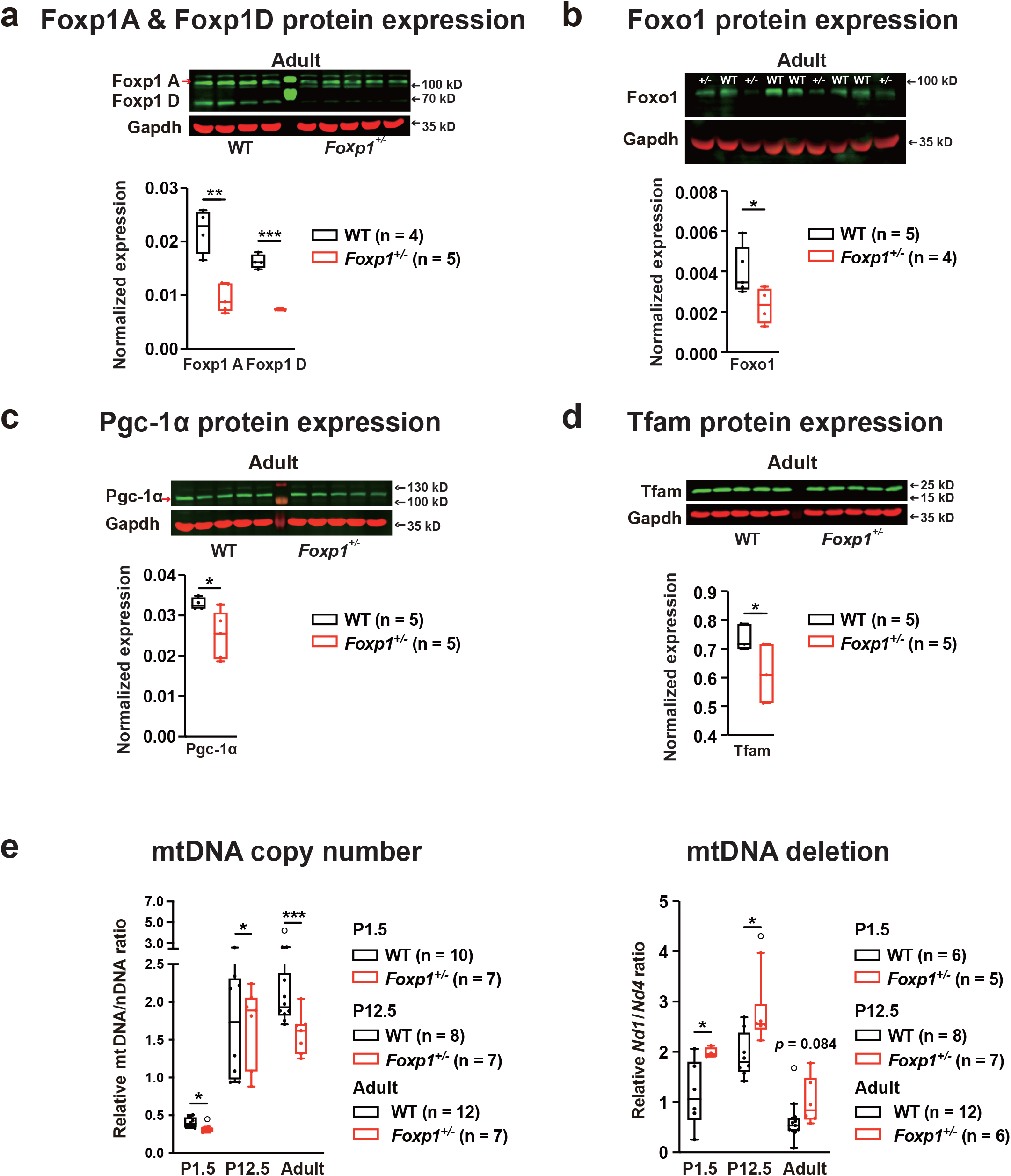
Mitochondrial biogenesis is altered in the *Foxp1*^+/−^ striatum. **a** The expression of Foxp1 isoform A and D were quantified in adult WT and *Foxp1*^+/−^ striatal tissue by western blot. At the age of 8 weeks Foxp1 A was reduced by ~57% and Foxp1 D by ~55% in *Foxp1*^+/−^ mice compared to WT animals. **b** Foxo1 protein was reduced by ~42% in adult *Foxp1*^+/−^ tissue, **c** Pgc-1α by ~24% and **d** Tfam by ~17%. **e** Mitochondrial copy number and mtDNA deletion were evaluated by quantitative real-time PCR. The mtDNA/nDNA ratio was calculated by normalizing the expression of *D-loop*, *16sRNA,* and *Nd1* to *Hk2* and *B2m*. mtDNA/nDNA ratios were reduced in *Foxp1*^+/−^ tissue at all three stages, indicating a decreased mitochondrial copy number. The *Nd1/Nd4* ratio was elevated in the *Foxp1*^+/−^ striatum at P1.5 and P12.5, indicating increased mtDNA deletions. In each experiment, at least four animals per group were used, the exact number of animals is given in the figure. In the box-and-whisker plot, the boxes represent the first and third quartiles, the whiskers represent 95% confidence intervals, and the lines within the boxes are median values. Weak outliers are marked with a circle. Asterisks indicate a significant difference (\**p* ≤ 0.05, \*\**p* ≤ 0.01, \*\*\**p* ≤ 0.001); two-way ANOVA.

The transcription factor Foxo1 is a “nutrient-sensor”. Depending upon the cellular stress levels, it shuttles between nucleus and mitochondria and binds to mitochondrial DNA (mtDNA) to regulate mitochondrial biogenesis and function [30, 31, 32]. Thus, reduced Foxo1 expression in *Foxp1*^*+/–*^ mice suggests that impaired mitochondrial biogenesis and function might play a role in the developmental defect of the *Foxp1*^+/−^ striatum [31, 32]. To test this hypothesis, we analyzed mRNA levels of *Pparα*, *Pparγ*, *Pgc-1α*, *Nrf1*, *Nrf2*, and *Tfam* at P1.5, P12.5, and 8 weeks, as they are known to be regulated by Foxo1 and are also involved in mitochondrial biogenesis [33, 34, 35, 36, 37]. The expression of all genes was significantly altered in the *Foxp1*^+/−^ striatum at one or more of the three stages investigated (Supplementary Fig. 1c). In particular, proliferator-activated receptor gamma coactivator 1-alpha (Pgc-1α) and mitochondrial transcription factor A (Tfam) were downregulated on protein level at all three stages in *Foxp1*^+/−^ striatal tissue (Fig. 1c, d and Supplementary Fig. 2c, d), which was not the case for nuclear respiratory factor 1 (Nrf1) (Supplementary Fig. 2e).

Next, we calculated mitochondrial DNA copy numbers and deletions using the mtDNA/nDNA and *Nd1*/*Nd4* ratio at P1.5, P12.5, and 8 weeks. The mtDNA/nDNA ratio was significantly reduced and the *Nd1/Nd4* ratio was significantly increased in *Foxp1*^+/−^ striata compared to WT striata (Fig. 1e), supporting the impairment of mitochondrial biogenesis in the *Foxp1*^+/−^ striatum.

### Genes involved in fission and fusion are dysregulated in the *Foxp1*^+/−^ striatum, pointing to impaired mitochondrial dynamics

We observed significantly reduced Pgc-1α expression in the *Foxp1*^+/−^ striatum (Fig. 1c). Pgc-1α regulates mitochondrial dynamics by influencing dynamin-like 120 kDa protein (*Opa1*), mitofusin-1 (*Mfn1*) and mitofusin-2 (*Mfn2*) – all involved in fusion – and dynamin-1-like protein (*Drp1*) and mitochondrial fission 1 protein (*Fis1*), which are involved in fission (Fig. 2a). Expression of these genes was also altered in the *Foxp1*^+/−^ striatum at one or more of the investigated time points (Supplementary Fig. 3). Mfn1 protein levels, were significantly reduced at P12.5, but elevated at 8 weeks in the *Foxp1*^+/−^ striatum compared to respective Mfn1 levels in WT (Fig. 2b, Supplementary Fig. 4a). Opa1 has two isoforms: L-Opa1 and S-Opa1. S-Opa1 protein expression was lower at all three time points. This increased the L-Opa1/S-Opa1 ratio (Fig. 2b, Supplementary Fig. 4b), suggesting disturbed maintenance of cristae structure in *Foxp1*^+/−^ striata [38]. Drp1 protein levels increased at P1.5 and decreased at P12.5 and 8 weeks in *Foxp1*^+/−^ striata (Fig. 2c, Supplementary Fig. 4c). Reversible Drp1 phosphorylation at Drp1(Ser637) blocks mitochondrial fission [39], so we measured Ser637-phosphorylated Drp1 and detected higher levels in the *Foxp1*^+/−^ striatum than in the WT striatum at all three stages (Fig. 2c, Supplementary Fig. 4c). Fis1 mediates fission by recruiting Drp1 to mitochondria and also promotes mitophagy. Significantly elevated Fis1 protein levels in *Foxp1*^+/−^ striata at all examined stages indicate increased mitochondrial fission (Fig. 2c, Supplementary Fig. 4d). However, Drp1 must be recruited from the cytoplasm to the mitochondrial surface to mediate the mitochondrial fission [40]. Higher Drp1 expression was detected in the cytosolic fraction at P12.5 and 8 weeks (Supplementary Fig. 4e), suggesting decreased fission in the *Foxp1*^+/−^ striatum.

**Fig. 2:**
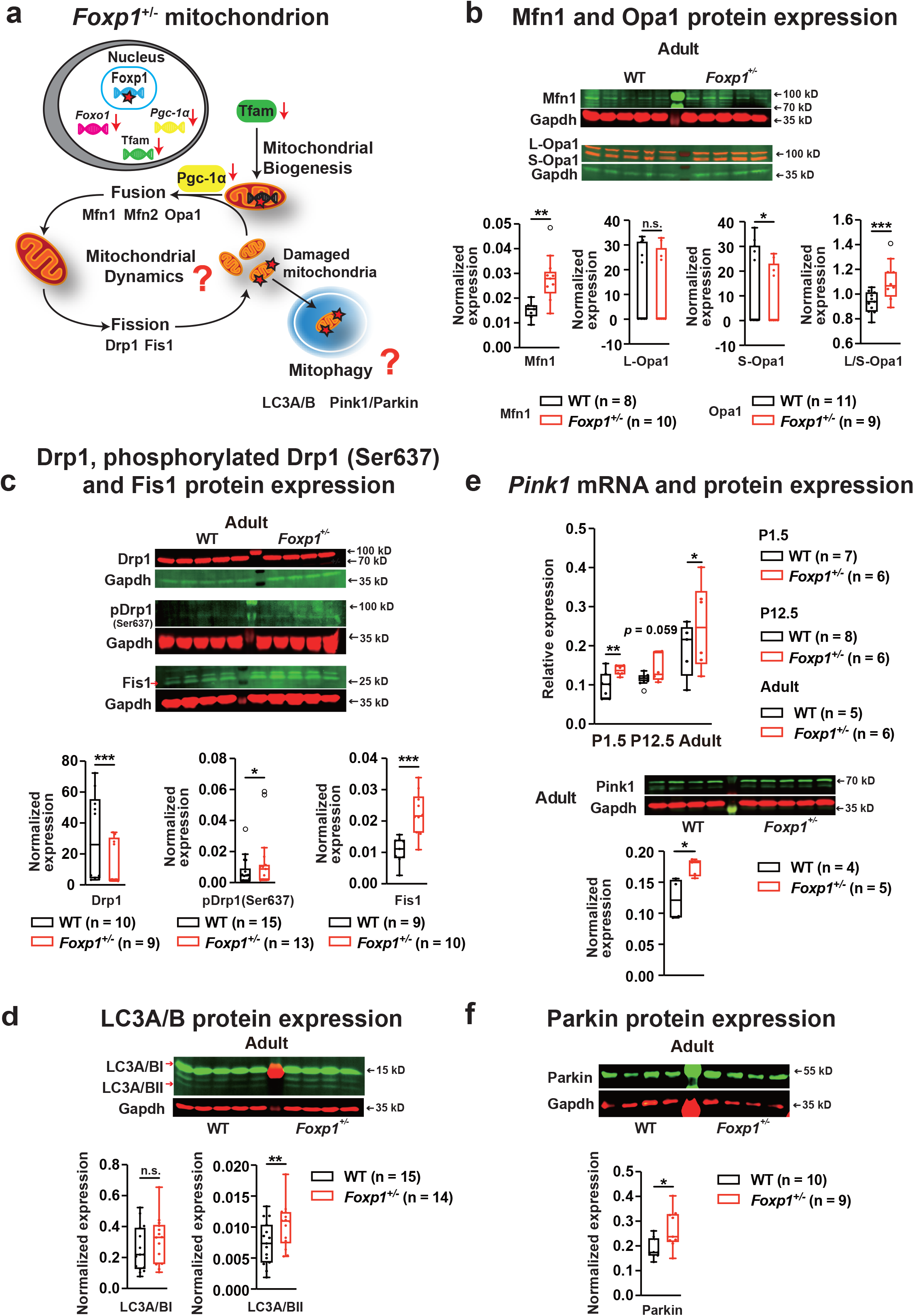
The *Foxp1*^+/−^ striatum displays altered mitochondrial dynamics and increased autophagy. **a** Scheme illustrating alterations of the mitochondrial machinery in the *Foxp1*^+/−^ striatum. Reduced Foxp1, Foxo1, Tfam, and Pgc-1α expression indicating disrupted mitochondrial biogenesis is represented by down-pointing arrows (red). Pgc-1α initiates mitochondrial dynamics by regulating genes involved in fusion (*Mfn1*, *Mfn2* and *Opa1*) and fission (*Drp1* and *Fis1*). LC3A/B and Pink-Parkin regulate mitophagy to eliminate damaged mitochondria. Mfn1, Opa1, Drp1, pDrp1 (Ser637), Fis1, LC3A/B Pink1 and Parkin were quantified by western blot at P1.5, P12.5 and adulthood (8 weeks). **b** Mfn1 expression was increased by ~89% in *Foxp1*^+/−^ striata at 8 weeks compared to WT tissue. Long (L)-Opa1 expression was not altered, short (S)-Opa1 expression by ~27% at 8 weeks in the *Foxp1*^+/−^ striatum. Consequently, the ratio of L-Opa1/S-Opa1 was significantly increased in the *Foxp1*^+/−^ striatum than in WT tissue at adult by ~20%. **c** Drp1 levels were increased by ~51% at 8 weeks in *Foxp1*^+/−^ animals. pDrp1 (Ser637) showed a significantly increased expression at adulthood compared to WT mice. Fis1 displayed an increased expression by ~109% at 8 weeks in the *Foxp1*^+/−^ striatum. **d** LC3A/BI and LC3A/BII isoform expression was quantified by western blot in striatal tissue of both genotypes. LC3A/BI expression did not differ between the genotypes, however, LC3A/BII levels were significantly increased in *Foxp1*^+/−^ tissue at 8 weeks. **e** *Pink1* mRNA levels were quantified by quantitative real-time PCR. *Pink1* expression levels were elevated by ~38% at P1.5 and by ~31% at the age of 8 weeks in *Foxp1*^+/−^ tissue. Pink1 expression was quantified by western blot and was increased in *Foxp1*^+/−^ tissue by ~42% at 8 weeks. **f** Parkin expression was quantified by western blot and was increased in *Foxp1*^+/−^ tissue by ~43% at 8 weeks. In each experiment, at least five animals per group were used, the exact number of animals is given in the figure. Weak outliers are marked with a circle. Asterisks indicate significant difference (\**p* ≤ 0.05, \*\**p* ≤ 0.01, \*\*\**p* ≤ 0.001); n.s. indicates non-significant; two-way ANOVA.

### Autophagy is increased in*Foxp1*^+/−^ striata

Damaged or excess mitochondria are eliminated by mitophagy. The microtubule-associated protein 1A/1B-light chain 3I (LC3I) is conjugated to phosphatidylethanolamine to form LC3II, a structural protein in autophagosomal membranes. We measured both LC3 isoform expressions to monitor autophagy and autophagic cell death. Western blot analysis showed that LC3A/BII expression was significantly higher in *Foxp1*^+/−^ than WT tissue at all three time points (Fig. 2d, Supplementary Fig. 5a). The Pink-Parkin pathway orchestrates the removal of damaged mitochondria in mitophagy. In line with this, we observed significantly elevated *Pink1* mRNA and its protein as well as Parkin protein expression in *Foxp1*^+/−^ striatal tissue, indicating increased autophagy (Fig. 2e-f, Supplementary Fig. 5b).

### Mitochondrial dysfunction and excessive oxidative stress in *Foxp1*^+/−^ striatal neurons

Proper mitochondrial dynamics, biogenesis, and mitophagy are essential for the maintenance of mitochondrial homeostasis and function. As mitochondria supply neurons with energy by generating metabolites and ATP via tricarboxylic acid cycle (TCA) and oxidative phosphorylation (OXPHOS), we measured adenosine compounds in primary striatal neurons. S-adenosylmethionine level was increased in *Foxp1*^+/−^ neurons, a sulfonium betaine that methylates DNA, RNA, and proteins in mitochondria and serves as an intermediate in the biosynthesis of lipoic acid and ubiquinone. However, the cellular content of ATP, ADP, AMP, NAD, NADH, 5‘-methylthioadenosine (MTA), adenosine and NADPH, did not significantly differ between WT and *Foxp1*^+/−^ neurons (Supplementary Fig. 6).

Mitochondrial membrane potential (Δψm) drives ATP generation, therefore we compared the Δψm in striatal neurons cultured from *Foxp1*^+/−^ and WT mice. The Δψm was lower in *Foxp1*^+/−^ neurons than in WT neurons, as shown by decreased Tetramethylrhodamine, methyl ester (TMRM; readily sequestered by active mitochondria) fluorescence, which indicate reduced OXPHOS in *Foxp1*^+/−^ mutant neurons (Fig. 3a). Thus, to directly estimate the rate of OXPHOS, we quantified Oxygen Consumption Rate (OCR), an indicator of the mitochondrial respiration, using the Seahorse metabolic flux analyzer (Supplementary Fig. 7a). Baseline mitochondrial respiration (Supplementary Fig. 7b) as well as the rate of OXPHOS (Supplementary Fig. 7c) were similar between *Foxp1*^+/−^ mutant and WT striatal neurons. In a pathological condition, to match the increased metabolic demands, cells tend to shift their metabolic state from OXPHOS towards aerobic glycolysis [41, 42]. Hence, to estimate whether such ‘metabolic state’ shift occurred in *Foxp1*^+/−^ mutant neurons, we quantified the rate of glycolysis by estimating Extracellular Acidification Rate (ECAR) in our seahorse metabolic flux analysis assays (Supplementary Fig. 7d). We found that *Foxp1*^+/−^ and WT striatal neurons performed glycolysis at similar level (Supplementary Fig. 7e).

**Fig. 3:**
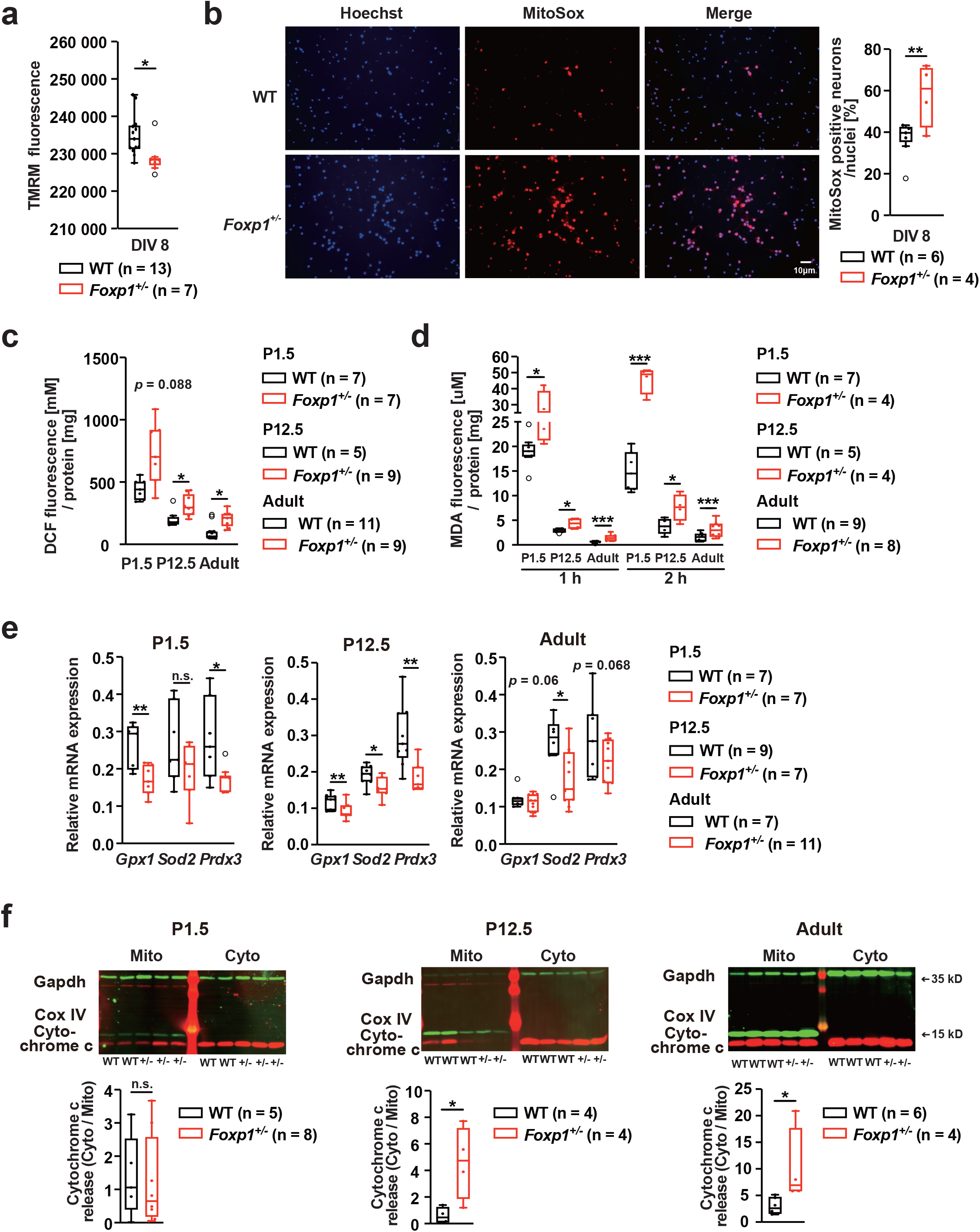
The *Foxp1*^+/−^ striatum exhibits mitochondrial dysfunction and increased oxidative stress. **a** Mitochondrial membrane potential (Δψm) was measured in primary striatal neurons of both genotypes by Tetramethylrhodamine, methyl ester (*TMRM*) staining. The TMRM fluorescent signal was significantly lower in *Foxp1*^+/−^ neurons compared to WT cells, indicating a lower mitochondrial membrane potential. **b** ROS abundance was quantified in living striatal neurons by MitoSOX™ Red, nuclei (blue) were stained with Hoechst 33324. *Foxp1*^+/−^ cultures had ~64% more MitoSOX™-red-positive neurons than WT cultures, indicating increased amounts of ROS. **c** DCF detection of striatal tissue at P1.5, P12.5, and 8 weeks. *Foxp1*^+/−^ tissue showed increased DCF fluorescence at P12.5 and 8 weeks compared to WT tissue indicating elevated ROS levels. **d** Measurement of lipid peroxidation in striatal tissue by malondialdehyde (MDA) fluorescence. At all three stages, MDA fluorescence was significantly increased in *Foxp1*^+/−^ tissue after 1 h and 2 h of incubation compared with WT tissue. **e** mRNA expression of *Gpx1*, *Sod2*, and *Prdx3* was analyzed by quantitative real-time PCR at P1.5, P12.5, and 8 weeks. *Foxp1*^+/−^ striata exhibited a reduced expression of *Gpx1* by ~35% and ~22% and of *Prdx3* by ~37% and ~38%, at P1.5 and P12.5, respectively. At P12.5 and 8 weeks of age *Sod2* levels were significantly decreased by ~12% and ~33 % compared to WT animals. **f** Quantification of cytochrome c in mitochondrial and cytosolic fractions of striatal tissue by western blot. Increased cytochrome c release in *Foxp1*^+/−^ animals indicated by increased cytosolic/mitochondrial cytochrome c ratio at P12.5 and 8 weeks by ~670% and~240%, respectively. In **a** and **b**, at least four animals per group were investigated and at least three replicates were performed per animal. The exact number of animals used is given in the figure. Cytosolic cytochrome c was normalized with Gapdh, mitochondrial cytochrome c with Cox IV. Weak outliers are marked with a circle. Asterisks indicate a significant difference (\**p* ≤ 0.05, \*\**p* ≤ 0.01, \*\*\**p* ≤ 0.001); n.s. indicates non-significant; two-way ANOVA. days *in vitro* (DIV).

Next, we quantified reactive oxygen species (ROS) in living primary striatal neurons and striatal tissue. Significantly more *Foxp1*^+/−^ neurons than WT neurons were stained positively with superoxide stain MitoSOX™ Red (Fig. 3b). More ROS were also detected in *Foxp1*^+/−^ striatal tissue than in WT striatal tissue as shown by quantification of 2’, 7’-dichlorofluorescein (DCF) (Fig. 3c). Moreover, lipid peroxidation was significantly higher in *Foxp1*^+/−^ striata than WT striata as assessed by malondialdehyde (MDA) formation (Fig. 3d). To assess anti-oxidative capacity, we measured glutathione peroxidase 1 (*Gpx1)*, superoxide dismutase 2 (*Sod2)*, and thioredoxin-dependent peroxide reductase (*Prdx3)*. We detected reduced *Gpx1* and *Prdx3* expression at P1.5 and P12.5 and reduced *Sod2* levels at P12.5 and 8 weeks in the *Foxp1*^+/−^ striatum (Fig. 3e). Elevated ROS at P12.5 and 8 weeks in *Foxp1*^+/−^ tissue suggests that *Sod2* might play the major role in ROS accumulation. Increased oxidative stress triggers apoptosis [43, 44]. To investigate apoptosis, we measured mitochondrial cytochrome c release, which is linked to early stages of apoptosis. Comparable to the analysis of ROS, cytochrome c levels in the cytosolic fraction did not differ at P1.5, but were significantly elevated in *Foxp1*^+/−^ tissue at P12.5 and 8 weeks (Fig. 3f). Therefore, we suggest that *Foxp1*^+/−^ mutant neurons exhibit increased mitochondrial ROS generation and at the same time, due to a reduced level of cellular anti-oxidants, these cells cannot optimally quench the detrimental effects of excessive ROS. Taken together, these results suggest *Foxp1*^+/−^ mutant neurons are able to maintain their metabolic state without switching between OXPHOS and glycolysis, but at a price of increased mitochondrial stress and ROS generation.

### Mitochondrial structural dynamics and dendritic arborization are altered in *Foxp1*^+/−^ striatal neurons

Mitochondrial stress and excessive ROS generation can lead to breakdown of mitochondrial network structure and influence the mitochondrial motility. To evaluate mitochondrial structure and dynamics, we stained neurons of both genotypes by using Mitotracker and performed live-cell 2-photon microscopy. Semi-automated segmentation of the mitochondria revealed that *Foxp1*^+/−^ neurons had a 19.2% decrease in mitochondrial surface area, compared to WT neurons (Fig. 4a). To track mitochondrial movement over time, we created kymographs using a semi-automatic algorithm in an image analysis software, ICY. *Foxp1*^+/−^ mitochondria displayed a consistent decrease in the total distance moved and the average speed of motility compared to WT mitochondria (Fig. 4b). We hypothesized that increased ROS and alterations in the mitochondrial network dynamics might impair neuronal morphology. Therefore, we examined the dendritic arbor complexity of GFP transfected WT and *Foxp1*^+/−^ striatal neurons by Sholl analysis after 8 days of differentiation, and found 45% fewer dendritic intersections in *Foxp1*^+/−^ neurons than in WT neurons (Fig. 4c). Hence, our results suggest that ongoing mitochondrial stress impairs mitochondrial structural dynamics and affects neuronal arborization.

**Fig. 4:**
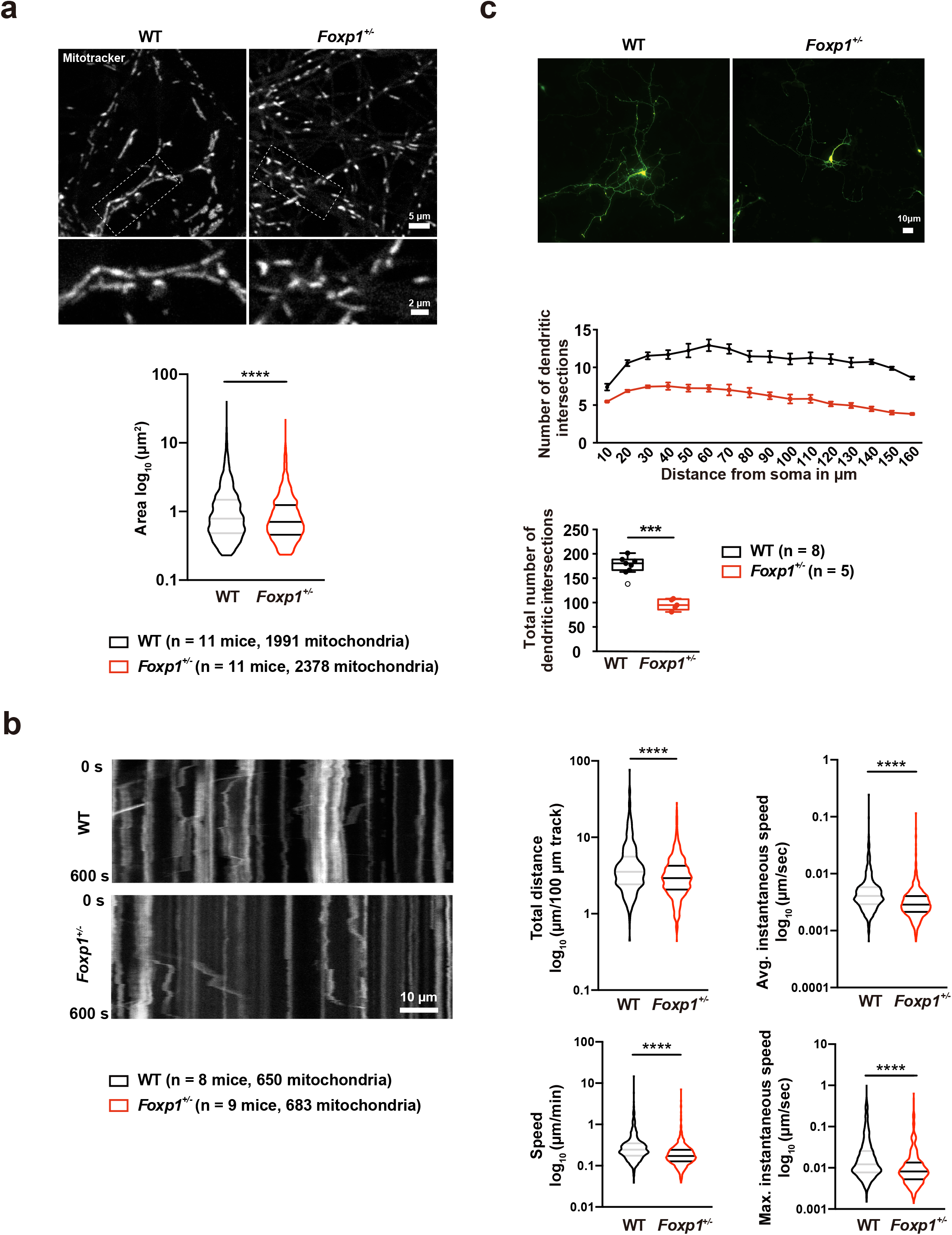
*Foxp1*^+/−^ striatal neurons show an altered mitochondrial structure and dynamics and dendritic arborization at DIV8. **a** Mitochondrial surface area was analyzed after staining with MitoTracker. There is no difference in the surface mitochondria content between genotypes (WT = 130.8 ± 23.17 μm^2^ (n = 11 mice, 20 images); *Foxp1*^+/−^ = 126.1 ± 19.92 μm^2^ (n = 11 mice, 20 images). *Foxp1*^+/−^ neurons displayed a decrease in mitochondrial surface area compared to WT cells (WT = 1.314 ± 1.777 μm2; *Foxp1*^+/−^ = 1.061 ± 1.202 μm^2^). 1991 WT and 2378 *Foxp1*^+/−^ mitochondria from 11 animals each were analyzed. **b** Kymographs of mitochondria in neurite extensions (10 min) demonstrate altered mitochondrial movement in *Foxp1*^+/−^ neurons. The distance of mitochondrial movement in *Foxp1*^+/−^ striatal neurons was reduced compared to WT mitochondria (WT 5.141 ± 5.823 μm/100 μm kymograph; *Foxp1*^+/−^ 3.568 ± 2.693 μm/100 μm kymograph). In addition, *Foxp1*^+/−^ mitochondria displayed a reduced speed as demonstrated by decreased displacement average speed (WT 0.363 ± 0.692 μm/min; *Foxp1*^+/−^ 0.229 ± 0.362 μm/min), instantaneous average speed (WT 0.0059 ± 0.0113 μm/sec; *Foxp1*^+/−^ 0.0038 ± 0.0059 μm/sec), and maximum instantaneous speed (WT 0.032 ± 0.068 μm/sec; *Foxp1*^+/−^ 0.018 ± 0.045 μm/sec). 650 WT and 683 *Foxp1*^+/−^ mitochondria from 8 WT and 9 *Foxp1*^+/−^ animals were analyzed. **c** Dendritic branching was investigated in GFP transfected primary striatal neurons by Sholl analysis. *Foxp1*^+/−^ neurons show an altered morphology with significant reduction of dendritic intersections by ~45% compared with WT neurons. In each experiment, at least five animals per group were investigated and at least three replicates were performed per animal. The exact number of animals used is given in the figure. **a, b** lines within violin plots represent the median and quartiles. Weak outliers are marked with a circle. Asterisks indicate a significant difference (\**p* ≤ 0.05, \*\**p* ≤ 0.01, \*\*\**p* ≤ 0.001, \*\*\*\**p* ≤ 0.0001); Mann-Whitney test and two-way ANOVA was performed in the analysis of mitochondrial dynamics and Sholl analysis. Days *in vitro,* DIV.

### *Foxp1*^+/−^ mice exhibit deficits in motor performance and in learning and memory

Striatal dysfunction probably contributes to the pathology of FOXP1 deficiency in a fundamental way. Since the striatum plays an essential role in motor control and motor learning, we investigated gait, coordination and endurance in *Foxp1*^+/−^ mice. The CatWalk gait analysis revealed a slow, hesitant walking pattern together with an increased number of steps, altered step characteristics and gait variability as well as disrupted interpaw coordination compared with WT mice (Fig. 5a and Supplementary Table 3). In the treadmill exhaustion test, *Foxp1*^+/−^ mice stopped running after 8 ± 2 min and covered a distance of 96 ± 37 m compared with WT mice, which stopped running after 16 ± 1 min and covered a distance of 245 ± 32 m. *Foxp1*^+/−^ mice also ran significantly slower than WT mice did (maximum speed 14 ± 2 m/min in *Foxp1*^+/−^ mice compared with 22 ± 1 m/min in WT mice) (Fig. 5b).

**Fig. 5:**
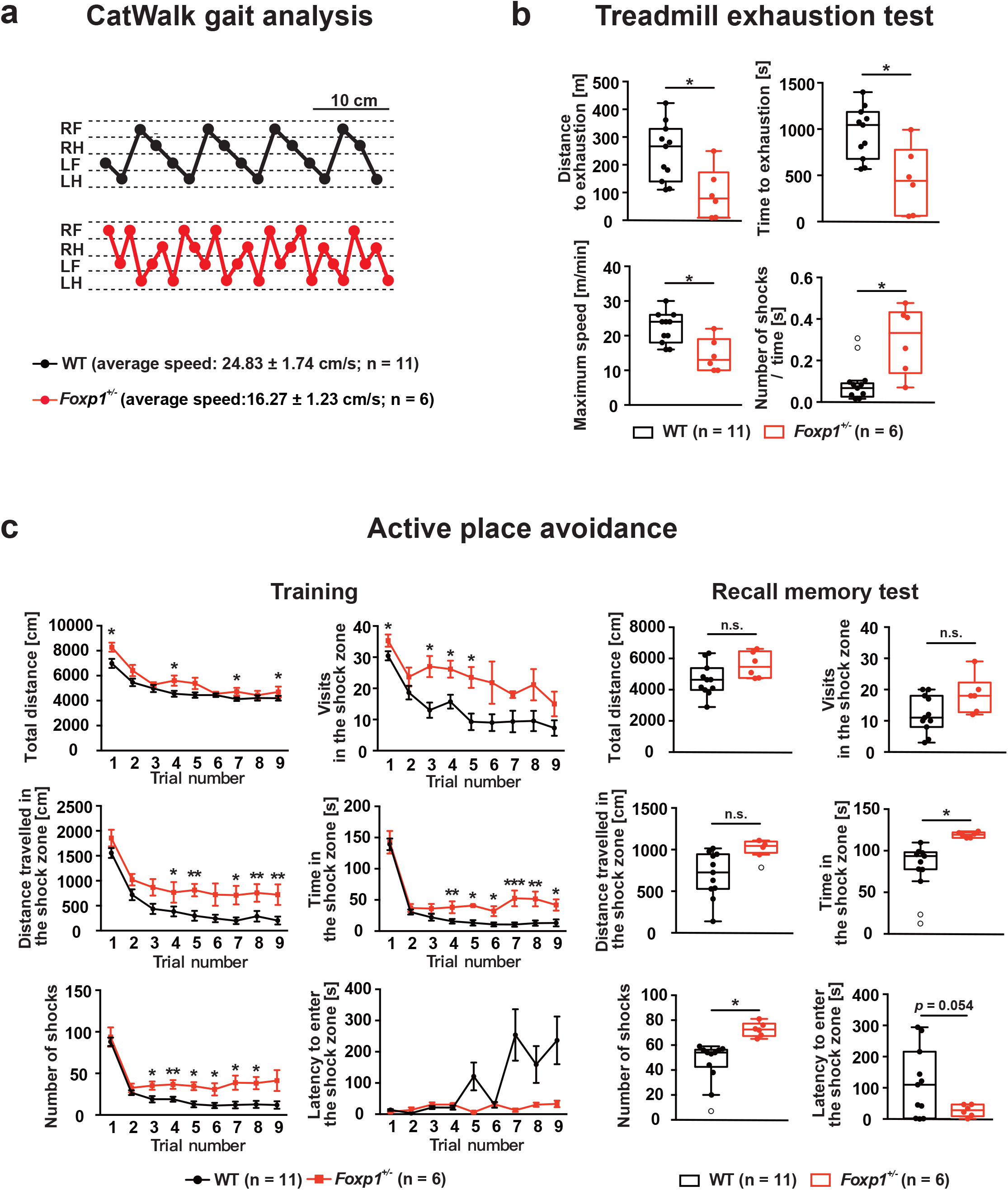
*Foxp1*^+/−^ mice show deficits in motor performance and cognition. **a** The walking pattern of WT and *Foxp1*^+/−^ mice was analyzed by the CatWalk gait analysis system. *Foxp1*^+/−^ mice exhibited a reduced walking speed by ~34% and their step number was significantly increased compared to WT mice. **b** Evaluation of endurance and muscle strength by treadmill exhaustion test. *Foxp1*^+/−^ mice covered a significantly decreased distance on the treadmill (96 ± 37 m versus 245 ± 32 m), exhausted earlier (8 ± 2 min versus 16 ± 1 min), exhibited a lower maximum speed (14 ± 2 m/min versus 22 ± 1 m/min), and received significantly more electrical shocks during the same period on the treadmill than WT animals. **c** Active place avoidance test. On day 1 in the training phase (9 trials), *Foxp1*^+/−^ mice displayed increased travelling and entered the shock zone more frequently than WT mice. In addition, they spent more time in there, covered a longer distance in the shock zone and thus received more shocks than WT mice. On day 2, when memory was tested, *Foxp1*^+/−^ mice entered the shock zone earlier, spent significantly more time in there, thus receiving ~60% more shocks than their WT littermates. In each experiment, at least six animals per group were used, the exact number of animals is given in the figure. In **c** course diagrams represent means ± SEM. Weak outliers are marked with a circle. Asterisks indicate significant difference (\**p* ≤ 0.05, \*\**p* ≤ 0.01, \*\*\**p* ≤ 0.001); n.s. indicates non-significant; three-way ANOVA. LF, left forelimb; RF, right forelimb; LH, left hindlimb; RH, right hindlimb.

We also tested active place avoidance because the striatum as a major component of the basal ganglia circuitry affects emotional and spatial learning [45, 46]. In this test, *Foxp1*^+/−^ mice could not avoid the shock zone. They entered the shock zone significantly more often, stayed there significantly longer, and covered a greater distance therein during the training phase, receiving more shocks than WT littermates did (Fig. 5c). The learning deficiency was confirmed in the following recall memory test. *Foxp1*^+/−^ mice spent more time in the shock zone and received more shocks than WT animals (Fig. 5c).

Together, we provide evidence of mitochondrial impairments and oxidative stress impacting on motor and cognitive deficits in *Foxp1*^+/−^ mice, which reflect core features of FOXP1 syndrome.

## Discussion

In this study, we show that *Foxp1* haploinsufficiency in mice leads to similar deficits in cognitive and motor function as reported in individuals with FOXP1 syndrome, and that mitochondrial dysfunction and oxidative stress is associated with these deficits.

Genes that regulate mitochondrial biogenesis and metabolism (*Foxo1*, *Pgc-1α*, *Nrf1*, *Nrf2*, *Tfam, Pparα*, and *Pparγ*) were all shown to be dysregulated in the *Foxp1*^+/−^ striatum. A comparison of these data to Nestin-Cre (*Foxp1*^-/-^) mice with a complete loss of *Foxp1* in the nervous system [12] and to mice with a complete loss of *Foxp1* in both D1-and D2-medium spiny neurons (MSNs) or D2-MSNs alone [15] revealed that reduced *Foxp1* expression in *Foxp1*^+/−^ mice affects gene expression differently than the complete loss of *Foxp1* does. An example is *Pgc-1α* [35, 47], regulated by Foxo1, which is a transcriptional coactivator and master regulator of mitochondrial biogenesis. Pparα and Pparγ interact with Pgc-1α [48, 49], suggesting that these proteins are part of a common pathway in the striatum. *Pgc-1α* has previously been shown to co-activate nuclear respiratory factor 2 (*Nrf2*), and together with Nrf2 coactivates *Nrf1* [50]. In turn, Pgc-1α and Nrf1/2 regulate *Tfam* transcription in response to increased mitochondrial biogenesis [51]. Decreased Foxo1 and Pgc-1α expression in the *Foxp1*^+/−^ striatum was detected by RNA-seq [9], confirming our data. The decreased Pgc-1α expression that we detected in the striatum is most likely caused by lowered Foxo1 activation, which may also explain the reduced *Nrf1* expression. Consequently, Pgc-1α deregulation may be responsible for the decreased expression of Tfam, which regulates mitochondrial genome replication and transcription [52, 53]. We also observed lower mtDNA/nDNA ratios, elevated *Nd1*/*Nd4* ratios, and elevated S-adenosyl methionine levels in the *Foxp1*^+/−^ striatum, indicating decreased mitochondrial copy numbers and increased mtDNA deletions [54].

In addition, we found clear evidence that proteins involved in fusion and fission are dysregulated in the mitochondria of the *Foxp1*^+/−^ striatum. Pgc-1α controls the delicate equilibrium between fusion and fission in mitochondria, and several genes/proteins associated with Pgc-1α were deregulated in the *Foxp1*^+/−^ striatum. Pgc-1α stimulates *Mfn1* [55], which, together with Mfn2, regulates the fusion of outer mitochondrial membranes. Interestingly, reduced Pgc-1α expression does not lead to significantly decrease Mfn1 mRNA and protein expression at P1.5. However, the adult *Foxp1*^+/−^ striatum reveals a reduced Mfn1 mRNA expression, whereas Mfn1 protein levels are significantly increased. This suggests a compensatory upregulation of Mfn1 translational. L- and S-Opa1 cooperate in mediating the fusion of inner mitochondrial membranes. L-Opa1 alone is sufficient for member docking and hemifusion, but shows a lower efficiency of content release by ~10% [56]. The pore opening efficiency of inner mitochondrial membrane can reach 80% upon addition of S-Opa1, which is fully competent for maintaining mitochondrial energetics and cristae structure [38, 56]. In the adult *Foxp1*^+/−^ striatum increased Mfn1 expression might enhance outer mitochondrial fusion to some extent. Nevertheless, we assume that reduced S-Opa1 may lead to inefficient and slow fusion of inner mitochondrial membrane, disrupted mitochondrial structure and energy supply. In addition, Drp1 (which activates fission) was reduced whereas phosphorylated-Ser637 Drp1 (which blocks fission) [39, 57] was increased in the *Foxp1*^+/−^ striatum, pointing to reduced mitochondrial fission. This finding is supported by elevated Drp1 levels in the cytosolic fraction of *Foxp1*^+/−^ tissue, hinting to more fission in the resting state.

Mitochondrial biogenesis and dynamics as well as the removal of damaged mitochondria are essential for mitochondrial function. Interestingly, we found that Fis1 expression is increased in the *Foxp1*^+/−^ striatum, and increased Fis1 expression has been associated with mitochondrial fragmentation and autophagosome formation [58]. LC3A/BII are microtubule-associated proteins that are essential for autophagosomes assembly [59]. Therefore, our finding that LC3A/BII expression is elevated in the *Foxp1*^+/−^ striatum indicates that autophagy is increased. We also found increased *Pink1* and Parkin expression in the *Foxp1*^+/−^ striatum, which may promote degradation of mitochondrial outer membrane proteins and autophagy of damaged mitochondria upon loss of mitochondrial membrane potential [60]. Strikingly, mitochondrial membrane potential was reduced in *Foxp1*^+/−^ striatal cells, which may influence ATP production. However, while ADP and AMP levels were increased compared with wild type, this was not the case for ATP levels, likely due to the high variance between striatal neurons.

We observed increased ROS production and lipid peroxidation as well as reduced expression of antioxidant genes and elevated cytochrome c release in *Foxp1*^+/−^ neurons and tissues, pointing to cellular injury and apoptosis [61]. Moreover, elevated ROS can in turn affect mitochondrial structure and motility [62, 63]. The significant reduction in mitochondrial surface area and average speed of *Foxp1*^+/−^ striatal neurons suggests an interaction between mitochondrial dysfunction and excessive ROS. In addition, mitochondrial dysfunction and increased ROS may affect cell morphology, and we observed a significantly lower number of dendritic intersections in *Foxp1*^+/−^ striatal neurons than in WT neurons. Altered dendritic growth and/or maintenance disrupt neural network function, ultimately leading to social and cognitive impairment. Indeed, impaired connectivity between the higher-order association areas is a major defect in ASD [64] and seems to hold true for FOXP1 syndrome as well.

Clinical studies have identified mitochondrial disturbances at the levels of mtDNA, complex I and III activity, oxidative stress, and metabolites in individuals with ASD [65, 66]. Mitochondrial dysfunction is found more frequently in individuals with ASD than in the general population, but seems to affect only a subset of individuals with ASD [19]. However, the ratio of mitochondrial dysfunction found in ASD individuals varies due to different selections of metabolic biomarkers (e.g. pyruvate, lactate, alanine) and biospecimens (e.g. blood, urine, muscle). A more accurate way to define this subgroup is the identification of dysregulated mitochondria-related genes. Several genes (e.g. *SLC25A12*, *MTX2*, *NEF*, *SLC25A27*, *SHANK3*, *SCO2*) that play a role in mitochondrial function are associated with the pathophysiology of autism [67, 68, 69]. Foxp1 is expressed in many tissues apart from the brain, suggesting that the mitochondrial defects that we observed in striatal tissue may also be present in other tissues with FOXP1 deficiency. This corresponds with the understanding that ASD is not solely a central nervous system disorder but also causes muscular, gastrointestinal, endocrine, and immune deficiencies [19, 70]. FOXP1 deficiency, therefore, defines one subgroup of ASD individuals affected by mitochondrial dysfunction.

Mitochondrial dysfunction and increased oxidative stress can interplay and activate components of pathways regulating autophagy, apoptosis, and inflammation. These defects will inevitably affect all areas of cognition and motor function that rely on the basal ganglia circuitry. Moreover, mitochondrial function plays an important role in neurogenesis, synaptic plasticity, and neurotransmission [16, 17, 18]. Very interesting in this context is the finding that disrupted mitochondrial activity cause GABA sequestration in mitochondria, reducing GABAergic signaling in the striatum and resulting in social deficits [71]. In this respect, we conclude that mitochondrial dysfunction does not only contribute to the pathology of FOXP1 syndrome, but also provides the impetus for the development of this disorder.

Our finding that mitochondrial function is disrupted in the *Foxp1*^+/−^ striatum may also be relevant to Huntington’s disease (HD). This neurodegenerative disorder is characterized by the degeneration of GABAergic medium spiny neurons and mitochondrial dysfunction [72, 73], but the molecular mechanism by which mutant Huntingtin affects mitochondrial function remains elusive. Strikingly, FOXP1 is downregulated in HD individuals and mouse models of HD [74, 75]. Elevating Foxp1 expression protects striatal and cortical neurons from death caused by mutant Huntingtin. Conversely, Foxp1 knockdown promotes death in healthy neurons, indicating a neuroprotective function for Foxp1 [74]. In addition, PGC-1α is downregulated in animal models of HD and individuals with HD, providing a possible link between impaired energy metabolism and neurodegeneration [76]. Taken together, these results suggest that mitochondrial dysfunction and energy homeostasis in HD may be caused by reduced FOXP1 expression.

The striatum plays a vital role in the coordination of motor and cognitive information from diverse brain regions into meaningful behavioral output [15]. Approximately 95% of the striatal cells are Foxp1-expressing medium spiny neurons, which, at least *in vitro* upon current injection spike at a higher firing rate (50 to 60 Hz) than pyramidal neurons [77, 78]. Whether such high firing rates are reached *in vivo* during certain behaviors is not known. Under standard recording conditions *in vivo*, medium spiny neurons exhibit sparse firing, but up-states and subthreshold depolarization may well contribute to significant calcium influx [79]. Thus, it may well be that these properties make medium spiny neurons prone to a higher energy demand and neuronal vulnerability. Considering that the striatum is strongly affected by Foxp1 deficiency, striatal dysfunction presumably plays a key role in the pathogenesis of FOXP1 syndrome.

Motor performance depends on proper muscle function, which is controlled by the striatum and spinal motor neurons. We have previously shown that conditional deletion of *Foxp1* in the central and peripheral nervous system leads to severe striatal degeneration [12]; Foxp1 also plays a crucial role in the columnar organization of spinal motor neurons [80]. Furthermore, muscular malfunction leads to achalasia and altered colon motility in *Foxp1*^+/−^ mice [10] and gastrointestinal problems in patients [2]. Our report on reduced muscle strength as well as disrupted gait and coordination in *Foxp1*^+/−^ mice, pointing to hypotonia and motor dysfunction, closely mirrors the phenotype of individuals with FOXP1 deficiency [11]. Impaired cognitive function is one of core features in FOXP1 syndrome. It was recently shown that output of the basal ganglia via the substantia nigra *pars reticulata* (SNr) gates active avoidance [46], and that the striatum plays an important role in this by regulating SNr cell activity. The complete loss of Foxp1 in striatum leads to severe morphological and functional disruptions [12]. Apart from the striatum, cognitive function is also depended on the proper expression of Foxp1 in the pyramidal neurons of the neocortex and the CA1/CA2 subfields of the hippocampus [13, 14]. Therefore, both alterations in the basal ganglia and corticohippocampal circuitry probably contribute to the deficits in learning and memory that we report here in *Foxp1*^+/−^ mice.

In conclusion, mitochondrial dysfunction and oxidative stress in the *Foxp1*^+/−^ striatum may explain the behavioral and motor deficits seen in mice and patients and provide an insight into the pathomechanisms underlying FOXP1 syndrome. Further studies, using cell-type and region-specific deletion or replenishment of Foxp1 are needed to determine whether defects in FOXP1 syndrome are restricted to the striatum or whether mitochondrial dysfunctions in other tissues contribute to its pathology. Current medical interventions for FOXP1 syndrome and autism focus on the behavioral phenotype. Strategies targeting electron transport chain, antioxidants, or the gut microbiome[81], however, might be more efficient therapies for FOXP1 syndrome. Thus, our study may offer a starting point for the development of new treatment options for this disorder.

## Acknowledgments

We acknowledge the expert advice of Dr. Claudia Pitzer and Barbara Kurpiers from the Interdisciplinary Neurobehavioral Core Unit, and thank the Metabolomics Core Technology Platform of the Excellence cluster “CellNetworks” (University of Heidelberg) (DFG grant ZUK 40/2010-3009262) for support. We also thank Prof. Dr. Stephan Herzig at the Joint UKHD-IDC Translational Diabetes Program for the use of their Seahorse Analyzer. The data storage service SDS@hd, supported by the Ministry of Science, Research and the Arts Baden-Württemberg and the DFG through the Grant INST 35/1314-1 FUGG are gratefully acknowledged. We are grateful to Dr. Rolf Sprengel, Institute for Anatomy and Cell Biology and Max Planck Institute for Medical Research, Heidelberg, Dr. Yu-Chao and Prof. Hannah Monyer, Department Clinical Neurobiology, Heidelberg as well as Dr. Yuting Ong, Prof. Beate Niesler, Dr. Claire Bacon and Beatrix Startt for helpful comments. We acknowledge the financial support from the China Scholarship Council (CSC; #201708230108) to J.W., the Chica and Heinz Schaller foundation to A.A., and the Humboldt Foundation to R.L.S. G.A.R. is a member of the CellNetworks Cluster of Excellence (EXC 81), Interdisciplinary Center for Neurosciences, and Center of Rare Disease.

## Author contributions

J.W. and H.F. conceived the study, designed experiments, and interpreted results. J.W. performed experiments, J.W. and H.F. collected, and analyzed the data. J.W. and H.F. wrote the draft. F.B. and R.L. performed the seahorse assay and mitotracker-staining experiments, A.A. helped to discuss the results. G.R. supervised the study and wrote the manuscript with comments from all authors.

## References

1. Horn D, Kapeller J, Rivera-Brugués N, Moog U, Lorenz-Depiereux B, Eck S, et al. Identification of FOXP1 deletions in three unrelated patients with mental retardation and significant speech and language deficits. Hum Mutat. 2010;31:E1851–60.

2. Siper PM, De Rubeis S, Trelles MDP, Durkin A, Di Marino D, Muratet F, et al. Prospective investigation of FOXP1 syndrome. Mol Autism. 2017;8:57.

3. Satterstrom FK, Kosmicki JA, Wang J, Breen MS, De Rubeis S, An JY, et al. Large-scale exome sequencing study implicates both developmental and functional changes in the neurobiology of autism. Cell. 2020;180:568–84

4. Li S, Weidenfeld J, Morrisey EE. Transcriptional and DNA binding activity of the Foxp1/2/4 family is modulated by heterotypic and homotypic protein interactions. Mol Cell Biol. 2004;24:809–22.

5. Mendoza E, Scharff C. Protein-protein interaction among the FoxP family members and their regulation of two target genes, *VLDLR* and *CNTNAP2* in the zebra finch song system. Front Mol Neurosci. 2017;10:112.

6. Lai CS, Fisher SE, Hurst JA, Vargha-Khadem F, Monaco AP. A forkhead-domain geneismutatedin a severe speech and language disorder. Nature. 2001;413:519–23.

7. Co M, Anderson AG, Konopka G. FOXP transcription factors in vertebrate brain development, function, and disorders. Wiley Interdiscip Rev Dev Biol. 2020;9:e375.

8. Wang B, Weidenfeld J, Lu MM, Maika S, Kuziel WA, Morrisey EE, et al. Foxp1 regulates cardiac outflow tract, endocardial cushion morphogenesis and myocyte proliferation and maturation. Development. 2004;131:4477–87.

9. Araujo DJ, Anderson AG, Berto S, Runnels W, Harper M, Ammanuel S, et al. FoxP1 orchestration of ASD-relevant signaling pathways in the striatum. Genes Dev. 2015;29:2081–96.

10. Fröhlich H, Kollmeyer ML, Linz VC, Stuhlinger M, Groneberg D, Reigl A, et al. Gastrointestinal dysfunction in autism displayed by altered motility and achalasia in *Foxp1*^+/−^ mice. Proc Natl Acad Sci USA. 2019;116:22237–45.

11. Meerschaut I, Rochefort D, Revençu N, Pètre J, Corsello C, Rouleau GA, et al. FOXP1-related intellectual disability syndrome: a recognisable entity. J Med Genet. 2017;54:613–23.

12. Bacon C, Schneider M, Le Magueresse C, Froehlich H, Sticht C, Gluch C, et al. Brain-specific *Foxp1* deletion impairs neuronal development and causes autistic-like behaviour. Mol Psychiatry. 2015;20:632–9.

13. Usui N, Araujo DJ, Kulkarni A, Co M, Ellegood J, Harper M, et al. Foxp1 regulation of neonatal vocalizations via cortical development. Genes Dev. 2017;31:2039–55.

14. Araujo DJ, Toriumi K, Escamilla CO, Kulkarni A, Anderson AG, Harper M, et al. Foxp1 in Forebrain Pyramidal Neurons Controls Gene Expression Required for Spatial Learning and Synaptic Plasticity. J Neurosci. 2017;37:10917–31.

15. Anderson AG, Kulkarni A, Harper M, Konopka G. Single-Cell Analysis of Foxp1-Driven Mechanisms Essential for Striatal Development. Cell Rep. 2020;30:3051–66 e7.

16. Kann O, Kovacs R. Mitochondria and neuronal activity. Am J Physiol Cell Physiol. 2007;292:C641–57.

17. Khacho M, Slack RS. Mitochondrial dynamics in the regulation of neurogenesis: From development to the adult brain. Dev Dyn. 2018;247:47–53.

18. Li Z, Okamoto K, Hayashi Y, Sheng M. The importance of dendritic mitochondria in the morphogenesis and plasticity of spines and synapses. Cell. 2004;119:873–87.

19. Rossignol DA, Frye RE. Mitochondrial dysfunction in autism spectrum disorders: a systematic review and meta-analysis. Mol Psychiatry. 2012;17:290–314.

20. Valiente-Pallejà A, Torrell H, Muntané G, Cortés MJ, Martínez-Leal R, Abasolo N, et al. Genetic and clinical evidence of mitochondrial dysfunction in autism spectrum disorder and intellectual disability. Hum Mol Genet. 2018;27:891–900.

21. Socci DJ, Bjugstad KB, Jones HC, Pattisapu JV, GW A. Evidence that oxidative stress is associated with the pathophysiology of inherited hydrocephalus in the H-Tx rat model. Exp Neurol. 1999;155:109–17.

22. Ferreira IL, Carmo C, Naia L, I. Mota S, Cristina Rego A. Assessing mitochondrial function in in vitro and ex vivo models of Huntington’s disease. Methods Mol Biol. 2018;1780:415–42.

23. Chenouard N, Buisson J, Bloch I, Bastin P, Olivo-Marin J-C. Curvelet analysis of kymograph for tracking bi-directional particles in fluorescence microscopy images. 2010 IEEE International Conference on Image Processing2010. p. 3657–60.

24. de Chaumont F, Dallongeville S, Chenouard N, Herve N, Pop S, Provoost T, et al. Icy: an open bioimage informatics platform for extended reproducible research. Nat Methods. 2012;9:690–6.

25. Berg S, Kutra D, Kroeger T, Straehle CN, Kausler BX, Haubold C, et al. ilastik: interactive machine learning for (bio)image analysis. Nat Methods. 2019;16:1226–32.

26. Le Fevre AK, Taylor S, Malek NH, Horn D, Carr CW, Abdul-Rahman OA, et al. FOXP1 mutations cause intellectual disability and a recognizable phenotype. Am J Med Genet A. 2013;161A:3166–75.

27. Krajeski RN, Macey-Dare A, van Heusden F, Ebrahimjee F, Ellender TJ. Dynamic postnatal development of the cellular and circuit properties of striatal D1 and D2 spiny projection neurons. J Physiol. 2019;597:5265–93.

28. Peixoto RT, Wang W, Croney DM, Kozorovitskiy Y, Sabatini BL. Early hyperactivity and precocious maturation of corticostriatal circuits in *Shank3B*-/-mice. Nat Neurosci. 2016;19:716–24.

29. Zou Y, Gong N, Cui Y, Wang X, Cui A, Chen Q, et al. Forkhead box P1 (FOXP1) transcription factor regulates hepatic glucose homeostasis. J Biol Chem. 2015;290:30607–15.

30. Lettieri-Barbato D, Ioannilli L, Aquilano K, Ciccarone F, Rosina M, Ciriolo MR. FoxO1 localizes to mitochondria of adipose tissue and is affected by nutrient stress. Metabolism. 2019;95:84–92.

31. Cheng Z, Guo S, Copps K, Dong X, Kollipara R, Rodgers JT, et al. Foxo1 integrates insulin signaling with mitochondrial function in the liver. Nat Med. 2009;15:1307–11.

32. Wang D, Wang Y, Zou X, Shi Y, Liu Q, Huyan T, et al. FOXO1 inhibition prevents renal ischemia-reperfusion injury via cAMP-response element binding protein/PPAR-gamma coactivator-1alpha-mediated mitochondrial biogenesis. Br J Pharmacol. 2020;177:432–48.

33. Bogacka I, Ukropcova B, McNeil M, Gimble JM, Smith SR. Structural and functional consequences of mitochondrial biogenesis in human adipocytes in vitro. J Clin Endocrinol Metab. 2005;90:6650–6.

34. Gan Z, Fu T, Kelly DP, Vega RB. Skeletal muscle mitochondrial remodeling in exercise and diseases. Cell Res. 2018;28:969–80.

35. Puigserver P, Rhee J, Donovan J, Walkey CJ, Yoon JC, Oriente F, et al. Insulin-regulated hepatic gluconeogenesis through FOXO1–PGC-1α interaction. Nature. 2003;423:550–55.

36. Sheng B, Wang X, Su B, Lee HG, Casadesus G, Perry G, et al. Impaired mitochondrial biogenesis contributes to mitochondrial dysfunction in Alzheimer’s disease. J Neurochem. 2012;120:419–29.

37. Ventura-Clapier R, Garnier A, Veksler V. Transcriptional control of mitochondrial biogenesis: the central role of PGC-1alpha. Cardiovasc Res. 2008;79:208–17.

38. Lee H, Smith SB, Yoon Y. The short variant of the mitochondrial dynamin OPA1 maintains mitochondrial energetics and cristae structure. J Biol Chem. 2017;292:7115–30.

39. Jahani-Asl A, Slack RS. The phosphorylation state of Drp1 determines cell fate. EMBO Rep. 2007;8:912–3.

40. Westermann B. Mitochondrial fusion and fission in cell life and death. Nat Rev Mol Cell Biol. 2010;11:872–84.

41. Koppenol WH, Bounds PL, Dang CV. Otto Warburg’s contributions to current concepts of cancer metabolism. Nat Rev Cancer. 2011;11:325–37.

42. Sun Y, Oravecz-Wilson K, Bridges S, McEachin R, Wu J, Kim SH, et al. miR-142 controls metabolic reprogramming that regulates dendritic cell activation. J Clin Invest. 2019;129:2029–42.

43. de Nigris F, Lerman A, Ignarro LJ, Williams-Ignarro S, Sica V, Baker AH, et al. Oxidation-sensitive mechanisms, vascular apoptosis and atherosclerosis. Trends in Molecular Medicine. 2003;9:351–59.

44. Mattson MP. Neuronal Life-and-Death Signaling, Apoptosis, and Neurodegenerative Disorders. ANTIOXIDANTS & REDOX SIGNALING. 2006;8:11 & 12.

45. Correia SS, McGrath AG, Lee A, Graybiel AM, Goosens KA. Amygdala-ventral striatum circuit activation decreases long-term fear. Elife. 2016;5:e12669.

46. Hormigo S, Vega-Flores G, Castro-Alamancos MA. Basal ganglia output controls active avoidance behavior. J Neurosci. 2016;36:10274–84.

47. Fernandez-Marcos PJ, Auwerx J. Regulation of PGC-1α, a nodal regulator of mitochondrial biogenesis. Am J Clin Nutr. 2011;93:884S–90.

48. Johri A, Chandra A, Flint Beal M. PGC-1α, mitochondrial dysfunction, and Huntington’s disease. Free Radic Biol Med. 2013;62:37–46.

49. Vega RB, Huss JM, Kelly DP. The coactivator PGC-1 cooperates with peroxisome proliferator-activated receptor α in transcriptional control of nuclear genes encoding mitochondrial fatty acid oxidation enzymes. Mol Cell Biol. 2000;20:1868–76.

50. Cartoni R, Léger B, Hock MB, Praz M, Crettenand A, Pich S, et al. Mitofusins 1/2 and ERRalpha expression are increased in human skeletal muscle after physical exercise. J Physiol. 2005;567:349–58.

51. Kang I, Chu CT, Kaufman BA. The mitochondrial transcription factor TFAM in neurodegeneration: emerging evidence and mechanisms. FEBS Lett. 2018;592:793–811.

52. Kelly DP, Scarpulla RC. Transcriptional regulatory circuits controlling mitochondrial biogenesis and function. Genes Dev. 2004;18:357–68.

53. Wu Z, Puigserver P, Andersson U, Zhang C, Adelmant G, Mootha V, et al. Mechanisms controlling mitochondrial biogenesis and respiration through the thermogenic coactivator PGC-1. Cell. 1999;98:115–24.

54. Lozoya OA, Martinez-Reyes I, Wang T, Grenet D, Bushel P, Li J, et al. Mitochondrial nicotinamide adenine dinucleotide reduced (NADH) oxidation links the tricarboxylic acid (TCA) cycle with methionine metabolism and nuclear DNA methylation. PLoS Biol. 2018;16:e2005707.

55. Martin OJ, Lai L, Soundarapandian MM, Leone TC, Zorzano A, Keller MP, et al. A role for peroxisome proliferator-activated receptor gamma coactivator-1 in the control of mitochondrial dynamics during postnatal cardiac growth. Circ Res. 2014;114:626–36.

56. Ge Y, Shi X, Boopathy S, McDonald J, Smith AW, Chao LH. Two forms of Opa1 cooperate to complete fusion of the mitochondrial inner-membrane. Elife. 2020;9:e50973.

57. Chang CR, Blackstone C. Cyclic AMP-dependent protein kinase phosphorylation of Drp1 regulates its GTPase activity and mitochondrial morphology. J Biol Chem. 2007;282:21583–7.

58. Gomes LC, Scorrano L. High levels of Fis1, a pro-fission mitochondrial protein, trigger autophagy. Biochim Biophys Acta. 2008;1777:860–6.

59. Tanida I, Ueno T, Kominami E. LC3 and Autophagy. Methods Mol Biol. 2008;445:77–88.

60. Pickles S, Vigié P, Youle RJ. Mitophagy and quality control mechanisms in mitochondrial maintenance. Curr Biol. 2018;28:R170–R85.

61. Chen Q, Chai YC, Mazumder S, Jiang C, Macklis RM, Chisolm GM, et al. The late increase in intracellular free radical oxygen species during apoptosis is associated with cytochrome c release, caspase activation, and mitochondrial dysfunction. cell death Differ. 2003;10:323–34.

62. Debattisti V, Gerencser AA, Saotome M, Das S, Hajnoczky G. ROS Control Mitochondrial Motility through p38 and the Motor Adaptor Miro/Trak. Cell Rep. 2017;21:1667–80.

63. Zorov DB, Juhaszova M, Sollott SJ. Mitochondrial reactive oxygen species (ROS) and ROS-induced ROS release. Physiol Rev. 2014;94:909–50.

64. Geschwind DH, Levitt P. Autism spectrum disorders: developmental disconnection syndromes. Curr Opin Neurobiol. 2007;17:103–11.

65. Giulivi C, Zhang YF, Omanska-Klusek A, Ross-Inta C, Wong S, Hertz-Picciotto I, et al. Mitochondrial dysfunction in autism. JAMA. 2010;304:2389–96.

66. Napoli E, Wong S, Giulivi C. Evidence of reactive oxygen species-mediated damage to mitochondrial DNA in children with typical autism. Mol Autism. 2013;4:2.

67. Anitha A, Nakamura K, Thanseem I, Yamada K, Iwayama Y, Toyota T, et al. Brain region-specific altered expression and association of mitochondria-related genes in autism. Mol Autism. 2012;3:12–12.

68. Frye RE, Cox D, Slattery J, Tippett M, Kahler S, Granpeesheh D, et al. Mitochondrial Dysfunction may explain symptom variation in Phelan-McDermid Syndrome. Sci Rep. 2016;6:19544.

69. Lepagnol-Bestel AM, Maussion G, Boda B, Cardona A, Iwayama Y, Delezoide AL, et al. SLC25A12 expression is associated with neurite outgrowth and is upregulated in the prefrontal cortex of autistic subjects. Mol Psychiatry. 2008;13:385–97.

70. Cheng N, Rho JM, Masino SA. Metabolic dysfunction underlying autism spectrum disorder and potential treatment approaches. Front Mol Neurosci. 2017;10:34.

71. Kanellopoulos AK, Mariano V, Spinazzi M, Woo YJ, McLean C, Pech U, et al. Aralar Sequesters GABA into Hyperactive Mitochondria, Causing Social Behavior Deficits. Cell. 2020;180:1178–97.

72. Costa V, Scorrano L. Shaping the role of mitochondria in the pathogenesis of Huntington’s disease. EMBO J. 2012;31:1853–64.

73. Ferrante RJ, Kowall NW, Richardson Jr. EP. Proliferative and degenerative changes in striatal spiny neurons in Huntington’s disease—a combined study using the section-golgi method and calbindin D28k immunocytochemistry. J Neurosci. 1991;11:3877–87.

74. Louis Sam Titus ASC, Yusuff T, Cassar M, Thomas E, Kretzschmar D, D’Mello SR. Reduced expression of Foxp1 as a contributing factor in Huntington’s disease. J Neurosci. 2017;37:6575–87.

75. Tang B, Becanovic K, Desplats PA, Spencer B, Hill AM, Connolly C, et al. Forkhead box protein P1 is a transcriptional repressor of immune signaling in the CNS: implications for transcriptional dysregulation in Huntington disease. Hum Mol Genet. 2012;21:3097–111.

76. McGill JK, Beal MF. PGC-1α, a new therapeutic target in Huntington’s disease? Cell. 2006;127:465–8.

77. Jonas P, Racca C, Sakmann B, Seeburg PH, Monyer H. Differences in Ca2+ Permeability of AMPA-type Glutamate Receptor Channels in Neocortical Neurons Caused by Differential GluR-B Subunit Expression. Neuron. 1994;12:1281–89.

78. Melzer S, Gil M, Koser DE, Michael M, Huang KW, Monyer H. Distinct Corticostriatal GABAergic Neurons Modulate Striatal Output Neurons and Motor Activity. Cell Rep. 2017;19:1045–55.

79. Carter AG, Sabatini BL. State-dependent calcium signaling in dendritic spines of striatal medium spiny neurons. Neuron. 2004;44:483–93.

80. Rousso DL, Gaber ZB, Wellik D, Morrisey EE, Novitch BG. Coordinated actions of the forkhead protein Foxp1 and Hox proteins in the columnar organization of spinal motor neurons. Neuron. 2008;59:226–40.

81. Castora FJ. Mitochondrial function and abnormalities implicated in the pathogenesis of ASD. Prog Neuropsychopharmacol Biol Psychiatry. 2019;92:83–108.

