## Supplementary figures, tables, materials and methods for "Mitochondrial dysfunction and oxidative stress contribute to cognitive and motor impairment in FOXP1 syndrome"

### Supplementary Information

#### CatWalk gait analysis

Gait patterns were analyzed with the CatWalk XT analysis system (Noldus version 10.6, Wageningen, Netherlands). This system consists of an enclosed walkway on a glass plate that is traversed by the mouse from one side of the walkway to the other. Green light which enters at the long edge of the plate is completely internally reflected. Where the paws of the animal touch the glass plate, light emerges leading to its scattering. The paws are captured by a high speed video camera that is positioned underneath the walkway by using Illuminated Footprint™ technology. The gait pattern was recorded according to pre-set paradigms [1].

#### Treadmill exhaustion test

A detailed protocol for treadmill is published elsewhere [2]. Briefly, the animals were forced to run on a treadmill (Shenyang Sino King Equipment, Shenyang, China) by light electric shocks. At first, the mice were trained for 5 min at a speed of 10 m/min for two consecutive days. On the third day, this initial speed was gradually increased by 2 m/min every 2 min, up to a maximum speed of 46 m/min. The experiment was stopped as soon as the mouse was exhausted and stayed in the shock zone for more than 12 s.

#### Active place avoidance (APA)

The APA is a circular metal arena shock grid underneath a rotating arena surrounded by a transparent wall (Sygnis Bioscience, Heidelberg, Germany). A random 60° sector was set as non-rotating shock zone, where the animals received a 0.4 mA electric shock upon entry and further identical shocks every 1.5 s, if they did not leave the sector. First, we let the animals explore the platform for 10 min; during this phase the shock zone was switched off. Then eight training rounds of 10 min each with the shock zone switched on were performed to train the animals, to avoid the shock sector. 24 h later the animals were tested once for 10 min, with the shock zone switched off. Mice were recorded and tracked during the entire experiment.

#### RNA isolation and cDNA synthesis

Total RNA was prepared from frozen mouse striatal tissue samples using peqGOLD TriFast™ (PEQLAB-Life Science, Radnor, Pennsylvania, USA). First strand cDNA synthesis was performed from 1.5 µg total RNA using a Superscript II reverse transcriptase kit (ThermoFisher Scientific, Waltham, MA, USA) and oligo dT12-18-primers (ThermoFisher Scientific, Waltham, MA, USA) according to the manufacturer’s instructions.

#### Quantitative real-time PCR

Quantitative real-time PCR was performed using the qTOWER system (Analytic Jena, Jena, Germany) with an annealing temperature of 60 °C using SYBR Green No-ROX Fast Mix (Bioline, Luckenwalde, Germany) according to the manufacturer's instructions. Each of the samples was analyzed in triplicate and relative mRNA levels were assessed using the Standard Curve Method by normalization to succinate dehydrogenase complex subunit A (*Sdha1*) and hypoxanthine phosphoribosyltransferase 1 (*Hprt1*). Primer sequences are listed in Supplementary Table 1.

#### Protein isolation and analysis

Total protein lysates were isolated from frozen mouse striatal tissue samples using RIPA buffer with cocktail protein inhibitor (ThermoFisher Scientific, Waltham, MA, USA). Proteins (30-60 µg) were separated on a 12% SDS-PAGE gel and analyzed by primary and secondary antibodies (Supplementary Table 2) according to manufacturer’s instructions. GAPDH was taken as reference using a primary mouse monoclonal/rabbit polyclonal antibody. IRDye 800CW and IRDye 680 (LICOR Biosciense, Lincoln, NE, USA) were used as secondary antibodies. Protein bands were visualized using the Odyssey Infrared Imaging System and quantified using Image Studio Lite 4.0 software (both LI-COR Biosciences, Lincoln, NE, USA).

#### Calculation of mitochondrial DNA copy number and mitochondrial DNA deletion

Mitochondrial DNA copy number was analyzed as previously described [3]. In brief, the expression of three mitochondrial genes *16sRNA, D-loop* and *Nd1* as well as the two nuclear genes *B2m* and *Hk2* was measured by quantitative real-time PCR and copy number was determined by the mtDNA/nDNA ratio. Evaluation of relative mitochondrial DNA deletion was performed as previously reported [4]. The assay is based on the measurement of two mitochondrial genes (*Nd1* and *Nd4*). In brief, we quantified a specific region within *Nd1*, which is rarely deleted in patients with mitochondrial diseases, and a region within *Nd4*, which was frequently found to be deleted in the majority of patients. An elevated *Nd1*/*Nd4* ratio indicates increased mtDNA deletion. Primer sequences are listed in Supplementary Table 1.

#### Isolation of mitochondria

Freshly dissected tissue was homogenized in ice-cold isolation buffer (20 mM HEPES, 250 mM sucrose, 10 mM KCl, 1.5 mM $MgCl_2$, 1 mM EGTA, 1 mM EDTA) supplemented with cocktail protease inhibitors (ThermoFisher Scientific, Waltham, MA, USA). After a differential centrifugation, the mitochondria-enriched pellet was cleaned with washing buffer (250 mM sucrose, 5 mM HEPES, 5 mM KOH, 0.1 mM EGTA, pH=7.2).

#### Primary striatal neuron culture and Sholl analysis

Striata were dissected from the brains of E14.5 embryos, mechanically dissociated by pipetting in neuron plating medium (DMEM, 10 % FCS) and plated onto Poly-L-Lysine (PLL) coated glass coverslips. After 24 h, medium was changed to neurobasal medium with B27 supplement. Every 48 h, half the medium was replaced with fresh neurobasal/B27 medium. After 7 days *in vitro* (DIV), striatal neurons were transfected with 1 µg eGFP construct using Lipofectamine, to visualize the entire neuronal morphology. Sholl analysis was performed as previously described [5]. Imaging was performed using a Nikon Ni-E microscope (Nikon Instruments Inc., Melville, NY, USA) and analyzed using Fiji (Java 8, National Institute of Health, USA).

#### Co-immunoprecipitation (Co-IP)

Co-immunoprecipatitation was carried out by using Dynabeads$^{TM}$ Protein G Immunoprecipitation Kit (ThermoFisher Scientific, Waltham, MA, USA) according to the manufacturer's instructions. Thereafter, proteins were analyzed by western blot. We used the following antibodies: Mouse IgG1 kappa isotype (negative control), Foxp1 and Foxo1 (listed in Supplementary Table 2).

#### Cellular adenosine compound measurements

Adenosine compounds were extracted from pellets of primary striatal neurons (DIV8) with 0.10 ml ice-cold 0.1 M HCl in an ultrasonic ice-bath for 10 min. The resulting homogenates were centrifuged for 10 min at 4°C and 16,400 g to remove cell debris. Adenosines were derivatized with chloroacetaldehyde as previously described [6] and separated by reversed phase chromatography on an Acquity HSS T3 column (100 mm x 2.1 mm, 1.7 µm, Waters) connected to an Acquity H-class UPLC system. Prior separation, the column was heated to 43 °C and equilibrated with 5 column volumes of buffer A (5.7 mM TBAS, 30.5 mM $KH_2PO_4$ pH 5.8) at a flow rate of 0.6 ml min$^{-1}$. Separation of adenosine derivates was achieved by increasing the concentration of buffer B (2/3 acetonitrile in 1/3 buffer A) in buffer A as follows: 1 min 1 % B, 2 min 8 % B, 3.2 min 14 % B, 9.5min 50 % B, and return to 1 % B in 1.5 min. The separated derivates were detected by fluorescence (Acquity FLR detector, Waters, excitation: 280 nm, emission: 410 nm) and quantified using ultrapure standards (Merck

KGaA, Darmstadt, Germany). Data acquisition and processing was performed with the Empower3 software suite (Waters, Milford, MA, USA).

#### Mitochondrial dynamics and structural analyses

Striatal neurons were seeded on Poly-L-Lysine (PLL) coated glass coverslips at 100.000 cells/□m$^2$ density, and maintained in growth medium until 8 DIV. Neurons were incubated with 100 nM MitoTracker Red CMXRos (Cat M7512. ThermoFisher) for 30 min at 37°C, and rinsed 3 times with growing medium. Live-cell imaging was performed using a custom built two-photon microscope (Bergamo II, Thorlabs, Newton, New Jersey, USA) using a Nikon NIR, 60X 1.0 NA objective. Two photon excitation was achieved using a mode-locked Ti: sapphire laser (Coherent Ultra II) tuned to 940 nm. Each image frame was acquired at the rate of 0.8 s/frame (about 1.25 Hz) for 600 seconds and was 111.15 µm × 111.15 µm at 1024 × 1024 pixel resolution. For the mitochondrial dynamics analyses, images were registered using ImageJ plugin Turboreg, and kymographs were generated on the time series images using the Icy Kymograph Tracker tool (version 2.1.2.0) [7, 8]. For mitochondrial structure analyses, Mitotracker-stained neurons were imaged and single frame images were segmented with Ilastik (version 1.4.0b7) [9], following the Pixel classification pipeline. Ilastik was trained for two labels (background and mitochondria), and the final segmented images were exported as binary masks. For mitochondria surface area measurements, binary masks were analyzed using the *Region Properties* MATLAB function (Mathworks, 2020a).

#### Oxygen Consumption Rate (OCR) and Extracellular Acidification Rate (ECAR)

Striatal neurons (45.000 cells/well) were seeded in Poly-L-Lysine (PLL) pre-coated Seahorse XF96 Cell Culture Microplate and cultured for 8 days. On the day of measurement, cells were washed 3 times and pre-incubated for 1 h in Seahorse XF96 Cell Culture medium (#103575-100) supplemented with 10 mM Glucose, 1 mM sodium pyruvate (Sigma, #S8636), 2 mM L-

Glutamine (Gibco, #A2916801). Oxygen Consumption Rate (OCR) and Extracellular Acidification Rate (ECAR) were measured by following XF Cell Mito Stress Test (Agilent User Guide 103015-100) and XF Glycolytic Rate Assay (Agilent User Guide 103344-100), respectively, according to the manufacturer's instructions. Briefly, OCR was measured under basal condition and after sequential injection of 1.5 μM oligomycin (Olig; Hello Bio, #HB4488), 0.5 μM ionophore 4-(trifluoromethoxy) phenylhydrazone (FCCP; Hello Bio, #HB2903), and 0.5 μM rotenone (Rot; Hello Bio, HB5398) plus 0.5 μM antimycin A (AntA; Sigma, #A8674). The following parameters were calculated according to the manufacturer's instructions, briefly described below:

- **Basal respiration:** value of last rate measurement before first injection – measurement rate after Rot/AntA injection (inhibitors of complexes III and I respectively);
- **ATP-linked respiration:** last rate measurement before Olig injection – minimum rate measurement after Olig injection (blocker of ATP-synthase);
- **Maximum respiratory capacity:** maximum rate measurement after FCCP injection (dissipator of proton gradient between the matrix and inner membrane space) measurement rate after Rot/AntA injection;
- **Proton leak:** minimum rate measurement after Olig injection – measurement rate after Rot/AntA injection.

ECAR was measured at basal condition and followed by sequential injection of 0.5 μM rotenone plus 0.5 μM antimycin A (inhibitors of complexes III and I, respectively) and injection of 10 mM 2-deoxyglucose (2DG; Sigma #D8375) (competitive inhibitor of hexokinase, the first enzyme in the glycolytic pathway). Values were normalized to cellular protein levels. Figures represent the values normalized by the basal measurement.

### Supplementary Figures

#### Supplementary Fig. 1: Reduced Foxp1 levels in the *Foxp1*$^{+/-}$ striatum lead to altered expression of genes involved in mitochondrial biogenesis

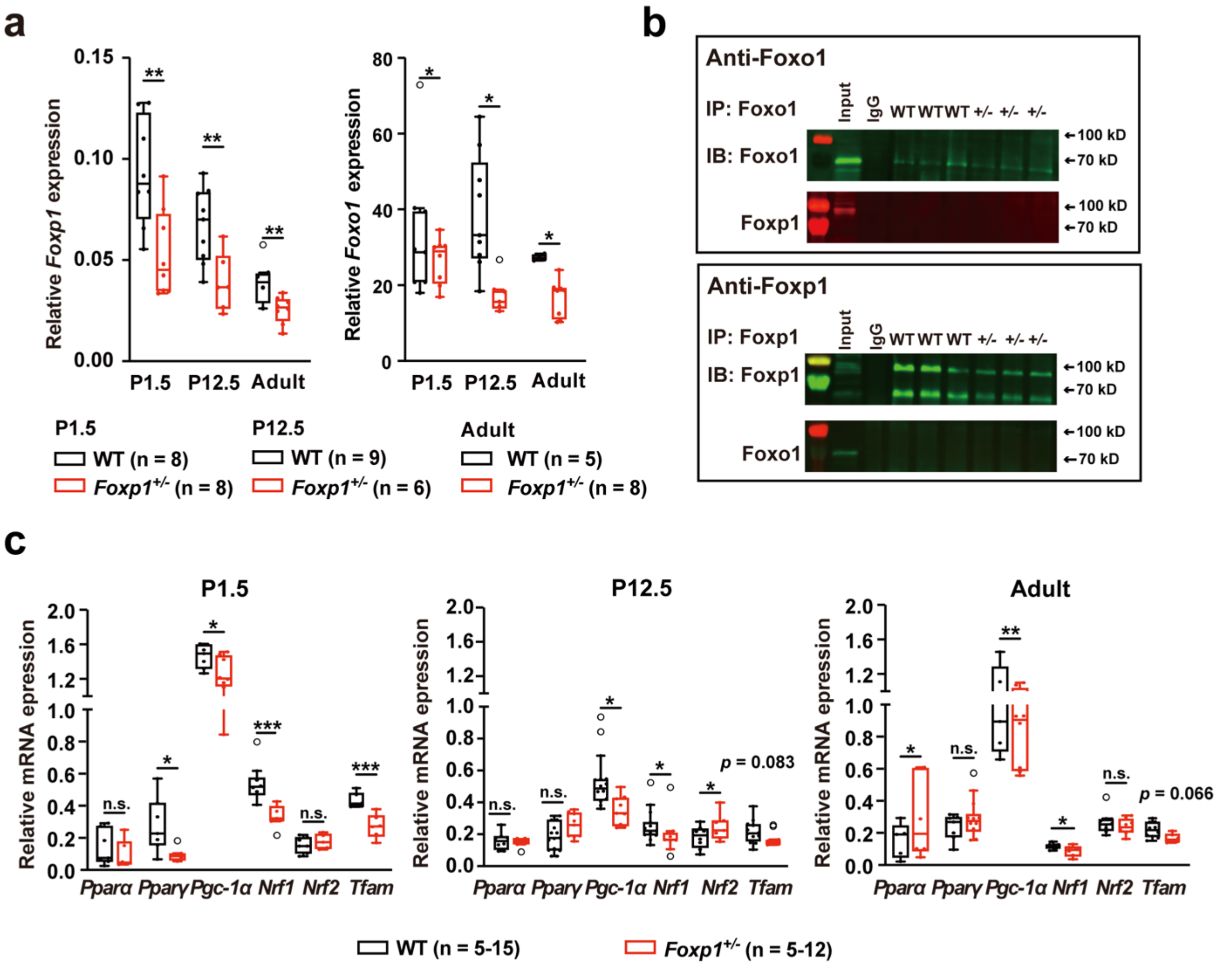


**a** *Foxp1* and *Foxo1* mRNA was quantified in WT and *Foxp1*$^{+/-}$ tissue at P1.5, P12.5 and 8 weeks by quantitative real-time PCR. *Foxp1* and *Foxo1* levels were significantly reduced in *Foxp1*$^{+/-}$ tissue on average by ~40% and ~35%, respectively. **b** Foxp1 and Foxo1 protein precipitation in WT and *Foxp1*$^{+/-}$ striatal tissue . Protein complexes were separated on SDS-PAGE and blotted for the indicated proteins. Mouse IgG kappa isotype was used as a negative control. No interaction between Foxp1 and Foxo1 was detected. **c** mRNA expression of *Pparα*, *Pparγ*, *Pgc-1α*, *Nrf1*, *Nrf2*, and *Tfam* was quantified in the WT and *Foxp1*$^{+/-}$ striatum at P1.5, P12.5 and 8 weeks by quantitative real-time PCR. All genes showed altered expression in *Foxp1*$^{+/-}$ tissue at at least one time point. On average, *Pgc-1α* and *Nrf1* were significantly downregulated by ~22% and ~33%, respectively. In each experiment, at least three animals per group were used, the exact number of animals is given in the figure. In the box-and-whisker plot, the boxes represent the first and third quartiles, the whiskers are 95% confidence interval, and the lines within the boxes are median values. Weak outliers are marked with a circle. Asterisks indicate a significant difference (*$p \leq 0.05$,**$p \leq 0.01$,***$p \leq 0.001$); n.s. represents non-significant; two-way ANOVA. Immunoprecipitation (IP), immunoblot (IB).

#### Supplementary Fig. 2: Proteins involved in mitochondrial biogenesis show an altered expression in the *Foxp1*$^{+/-}$ striatum

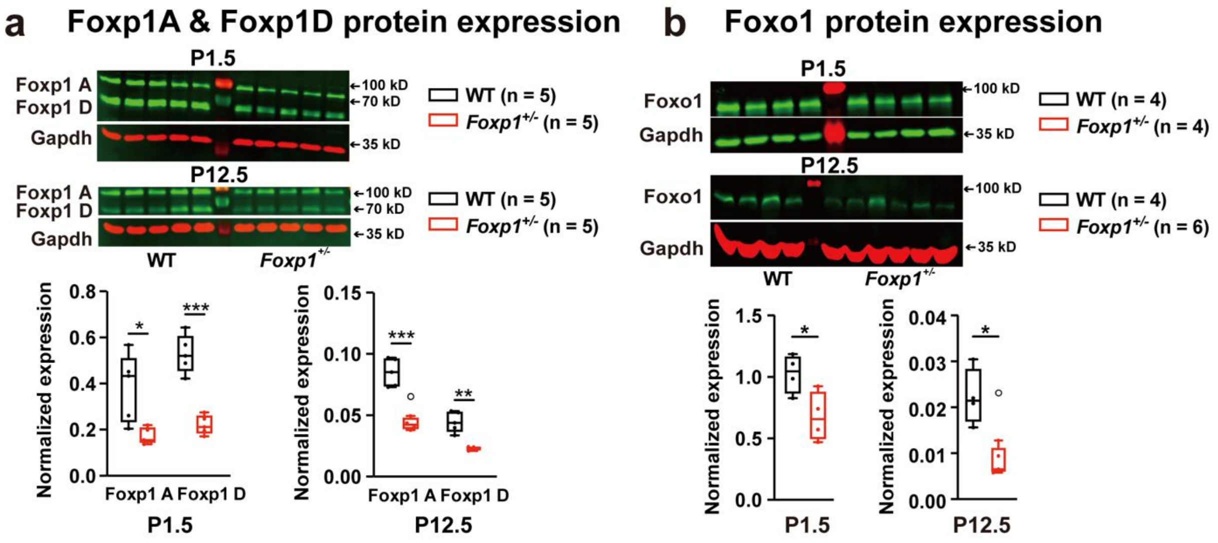

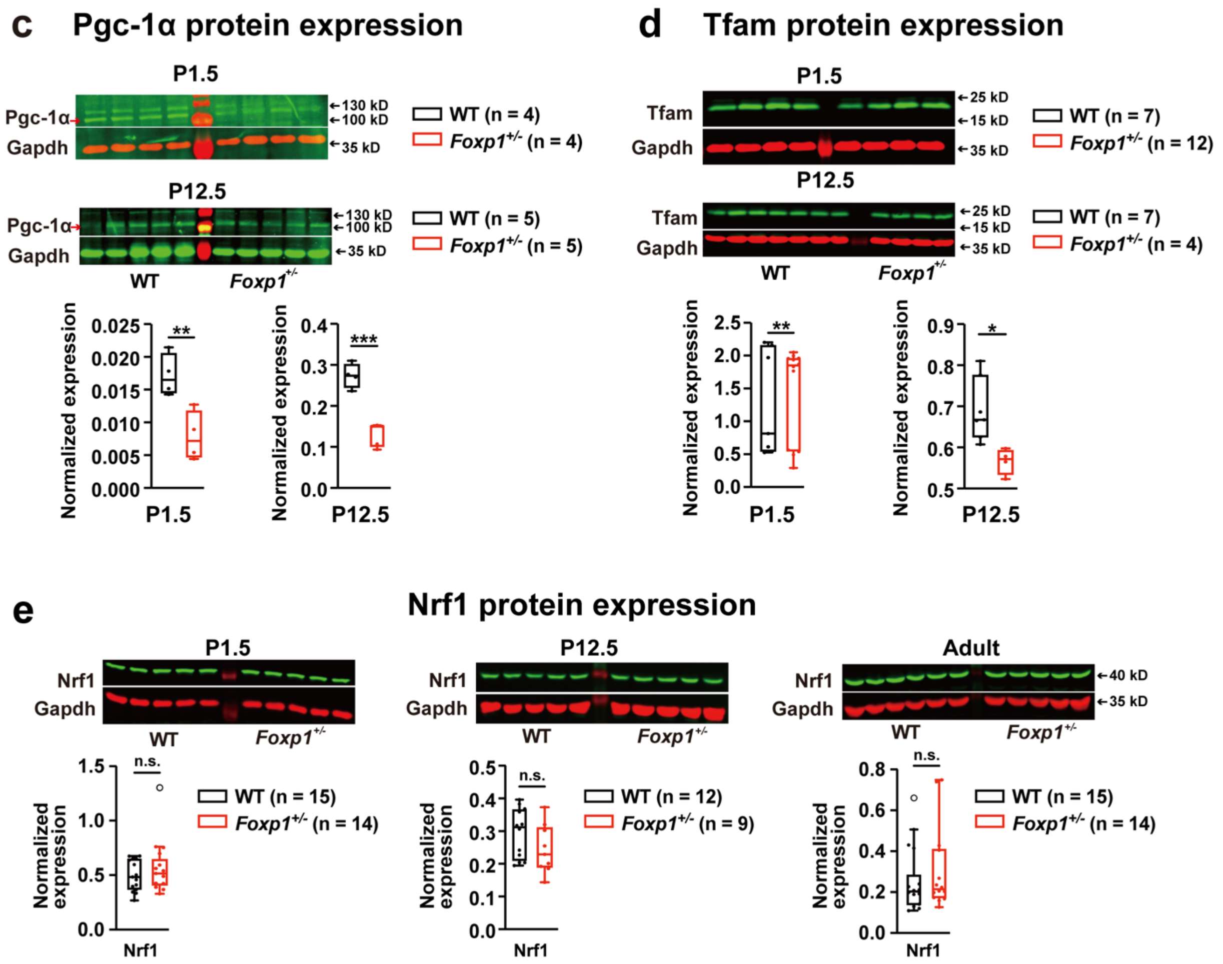


**a-d** Foxp1A, Foxp1D, Foxo1, Pgc-1α and Tfam levels were reduced in *Foxp1*$^{+/-}$ tissue at P1.5 by ~55%, ~58%, ~34%, ~54%, and ~12%, and at P12.5 by ~45%, ~43%, ~52%, ~52%, and ~18%, respectively. **e** Nrf1 expression did not differ between the genotypes at any of the three stages. In each experiment, at least four animals per group were used, the exact number of animals is given in the figure. Weak outliers are marked with a circle. Asterisks indicate a significant difference (**p* ≤ 0.05,***p* ≤ 0.01,****p* ≤ 0.001); n.s. represents non-significant.

**Supplementary Fig. 3**: **Genes involved in mitochondrial dynamics exhibit an altered expression in the *Foxp1*$^{+/-}$ striatum**

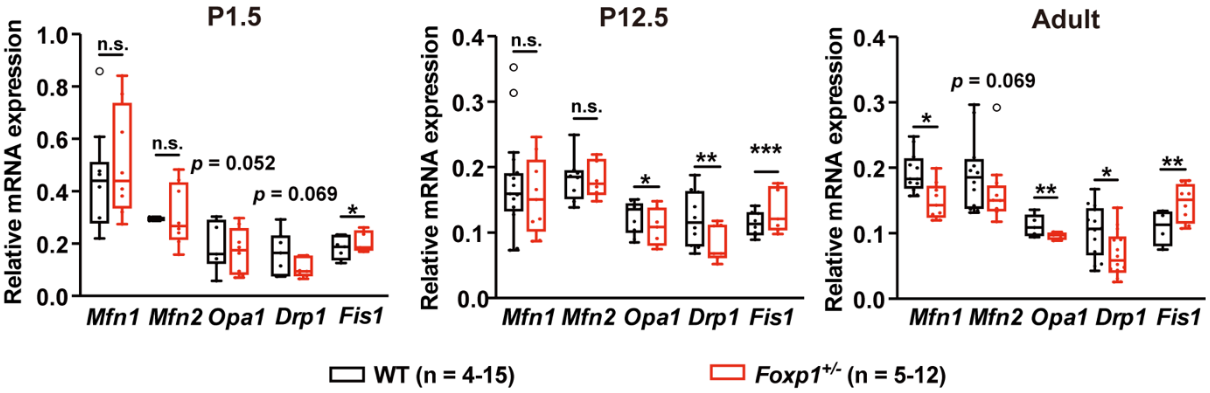


mRNA expression of *Mfn1*, *Mfn2*, *Opa1*, *Drp1*, and *Fis1* was measured by quantitative real-time PCR at P1.5, P12.5 and 8 weeks. Most genes showed an altered expression in at least one time point. *Fis1* expression was significantly increased at all three stages by ~13%, ~18% and ~34%, respectively. *Opa1* and *Drp1* were significantly decreased by ~13% and ~30% at P12.5, by ~13% and ~35% at 8 weeks. *Mfn1* was significantly reduced at 8 weeks, however, *Mfn2* was not altered at any of the three timepoints. In each experiment, at least four animals per group were used, the exact number of animals is given in the figure. Weak outliers are marked with a circle. Asterisks indicate significant difference (*$p \leq 0.05$, **$p \leq 0.01$, ***$p \leq 0.001$); n.s. represents non-significant; two-way ANOVA.

**Supplementary Fig. 4: Proteins involved in mitochondrial dynamics display an altered expression in the *Foxp1*$^{+/-}$ striatum at P1.5 and P12.5**

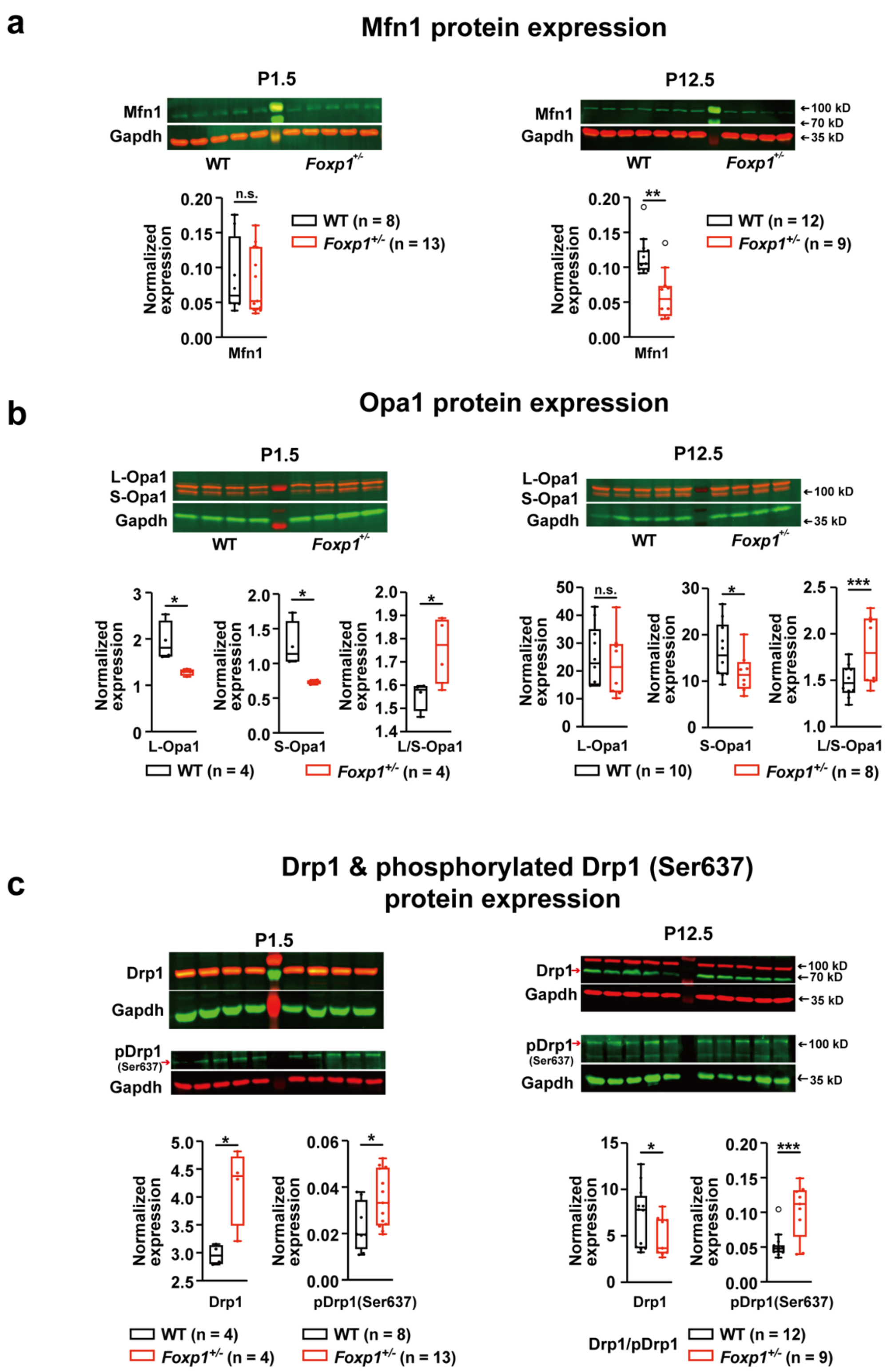

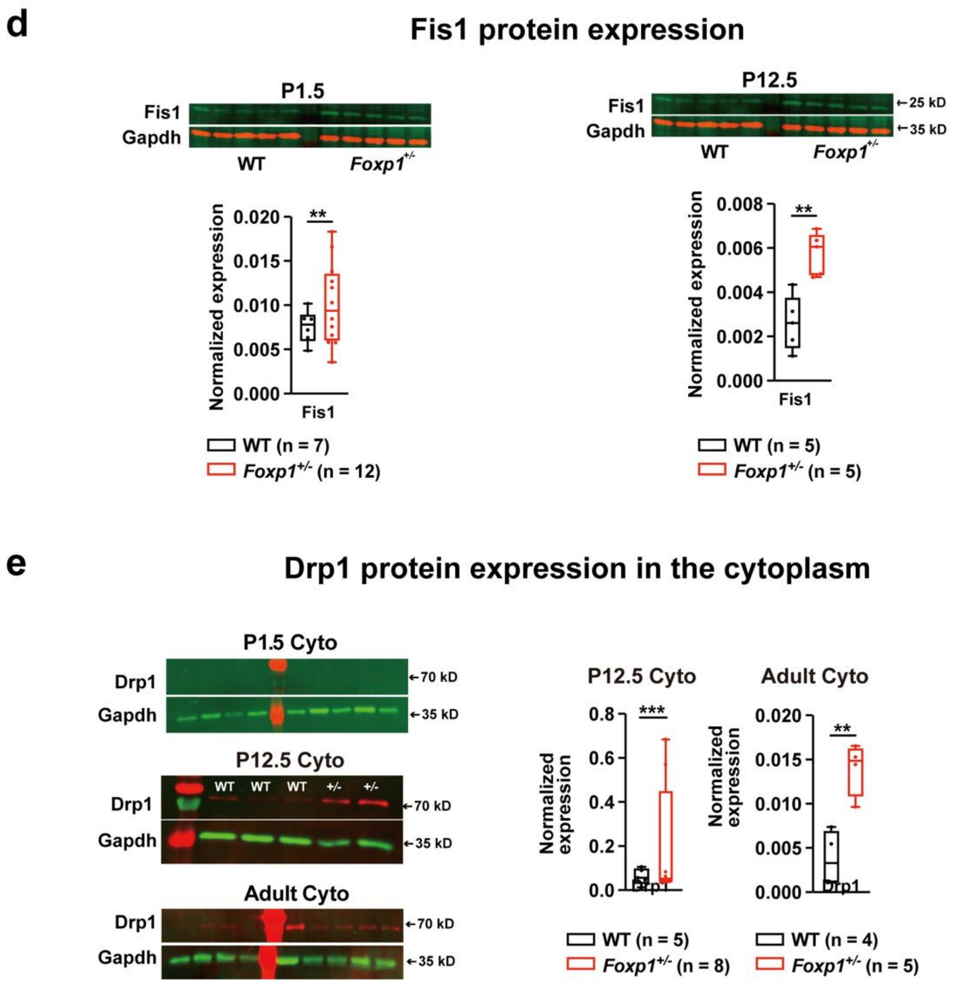


**a** Mfn1 expression showed no differences at P1.5, but it was reduced by ~45% at P12.5 in *Foxp1*$^{+/-}$ striata compared to WT tissue. **b** Long (L)-Opa1 expression was reduced by ~34% at P1.5, short (S)-Opa1 expression by ~42% at P1.5 and by ~29% at P12.5 in the *Foxp1*$^{+/-}$ striatum. The ratio of L-Opa1/S-Opa1 was significantly increased in the *Foxp1*$^{+/-}$ striatum than in WT tissue by ~13% at P1.5 and by ~22% at P12.5. **c** Drp1 levels were increased by ~42% at P1.5 and decreased by ~30% at P12.5 in *Foxp1*$^{+/-}$ animals whereas pDrp1 (Ser637) showed a significantly increased expression at P1.5 and P12.5 compared to WT mice. **d** Fis1 displayed an increased expression by ~34% at P1.5 and ~121% at P12.5 in the *Foxp1*$^{+/-}$ striatum. **e** Quantification of cytosolic Drp1. Drp1 was not detectable at P1.5 (n = 4 WT/5 *Foxp1*$^{+/-}$). At P12.5, *Foxp1*$^{+/-}$ tissue exhibited an increased Drp1 expression by ~214% and by ~342% at 8 weeks. In each experiment, at least four animals per group were used, the exact number of animals is given in the figure. Weak outliers are marked with a circle. Asterisks indicate significant difference (*$p$ ≤ 0.05, **$p$ ≤ 0.01, ***$p$ ≤ 0.001); n.s. represents non-significant; two-way ANOVA.

#### Supplementary Fig. 5: *Foxp1*$^{+/-}$ striatum shows increased expression of LC3A/BII and Parkin compared to WT striatum

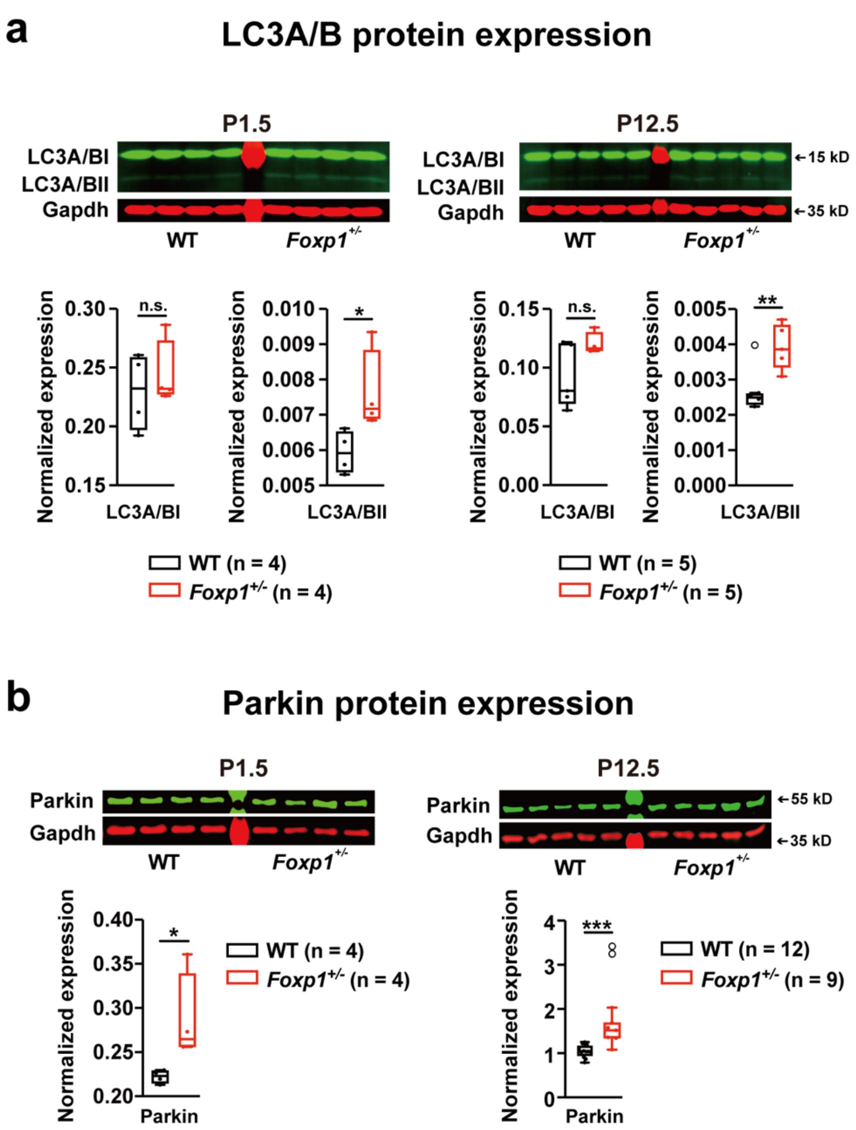


**a** LC3A/BI and LC3A/BII isoform expression was quantified by western blot in striatal tissue of both genotypes. LC3A/BI expression did not differ between the genotypes, however, LC3A/BII levels were significantly increased in *Foxp1*$^{+/-}$ tissue at P1.5 and P12.5. **b** Parkin expression was quantified by western blot and was increased in *Foxp1*$^{+/-}$ tissue by ~29% at

P1.5 and ~83% at P12.5. In each experiment, at least four animals per group were used, the exact number of animals is given in the figure. Weak outliers are marked with a circle. Asterisks indicate significant difference (*$p \leq 0.05$, **$p \leq 0.01$, ***$p \leq 0.001$); n.s. represents non-significant; two-way ANOVA.

#### Supplementary Fig. 6: *Foxp1*$^{+/-}$ primary striatal neurons display increased amounts of S-adenosyl methionine compared to WT cells at DIV8

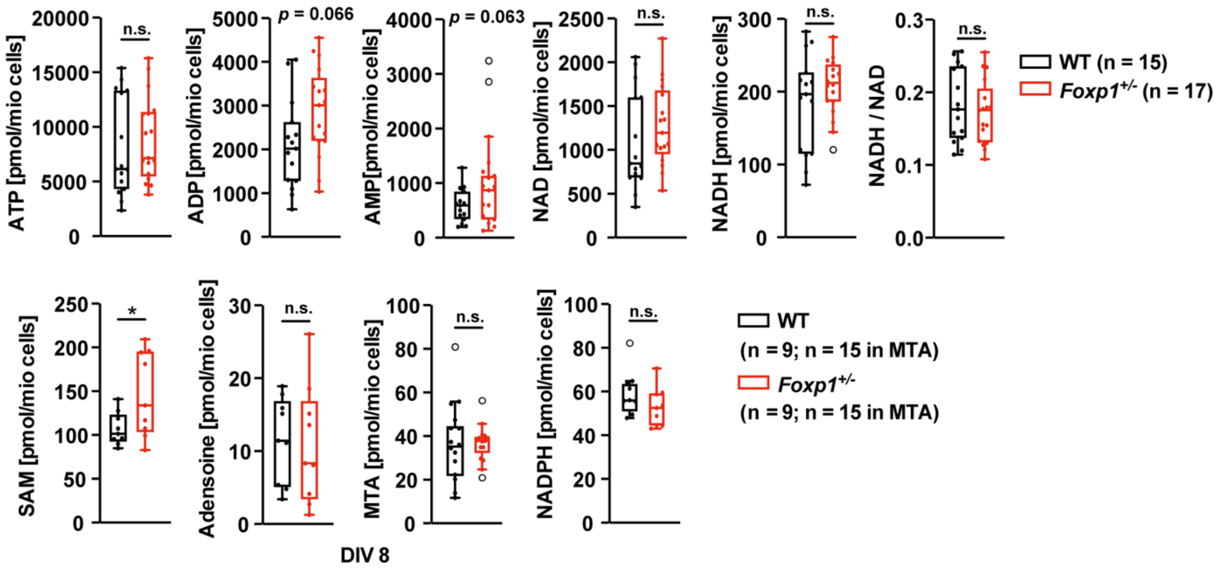


The mitochondrial metabolism associated products ATP, ADP, AMP, NAD, NADH, S-adenosyl methionine (SAM), adenosine, MTA and NADPH were quantified in WT and *Foxp1*$^{+/-}$ striatal neurons by UPLC. Except for increased SAM levels, *Foxp1*$^{+/-}$ neurons did not show significant differences compared with WT neurons. In each experiment, at least nine animals per group were used, the exact number of animals is given in the figure. Weak outliers are marked with a circle. Asterisks indicate significant difference (*$p \leq 0.05$); n.s. represents non-significant; two-way ANOVA. adenosine triphosphate (ATP); adenosine diphosphate (ADP); adenosine monophosphate (AMP); nicotinamide adenine dinucleotide (NAD); nicotinamide adenine dinucleotide-hydrogen (NADH); methylthioadenosine (MTA);

nicotinamide adenine dinucleotide phosphate (NADPH); ultra performance liquid chromatography (UPLC); days in vitro (DIV).

#### Supplementary Fig. 7: Oxygen consumption rate and extracellular acidification rate in striatal neurons at DIV8

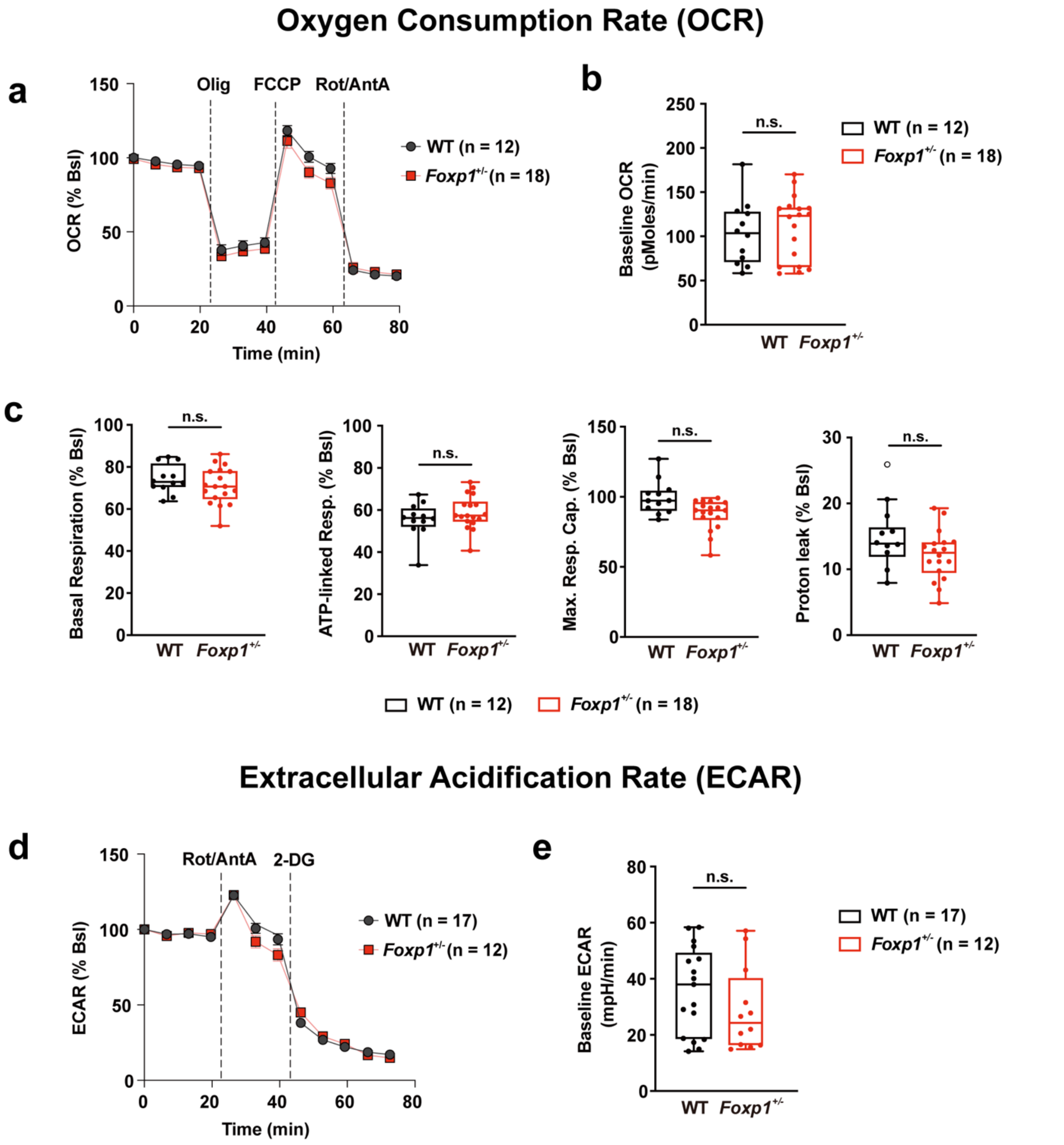

**a** Oxygen Consumption Rate (OCR) determined by Seahorse Bioscience XF96 analyzer in neurons treated sequentially with oligomycin (Olig), FCCP, and rotenone plus antimycin (Rot/Ant). OCR levels of each sample were normalized by the baseline (Bls). **b** Average of basal OCR values (baseline) in WT and *Foxp1*$^{+/-}$ neurons. **c** Basal respiration, ATP-linked respiration, maximum respiratory capacity and proton leak depicted for both genotypes. *Foxp1*$^{+/-}$ neurons exhibit a significantly reduced proton leak compared to WT neurons. **d** Extracellular Acidification Rate (ECAR) after sequential treatment with rotenone plus antimycin A antimycin (Rot/Ant) and 2-deoxyglucose (2-DG). ECAR levels were normalized by the baseline (Bls). **e** Average of basal ECAR levels (baseline) shown for both genotypes. In each experiment, at least 12 animals per group were investigated and at least three replicates were performed per animal. The exact number of animals used is given in the figure. Data represent the mean ± S.E.M. from biological replicates. Each biological replicate is the average of 2-6 technical replicates. Asterisks indicate significant difference (*$p \leq 0.05$) and n.s. indicates non-significant. Student t-test.

### Supplementary Tables

#### Supplementary Table 1: Primer list

| Genes | Sequence |
|---|---|
| *Foxp1 430bp 1* | 5'-CCTCTGGCGATGAACCTAGTGGT-3' |
| *Foxp1 430bp 2* | 5'-AGCCACACTTTCTCTCAGGATGT-3' |
| *Foxp1 280bp 1* | 5'-AGCGCATGCTCCAGACTGCCTTG-3' |
| *Foxp1* | For 5'-AGAGCGCCTGCAAGCCATGA-3' |
| | Rev 5'-GGCGGTGGGGGTTGTTGGAG-3' |
| *Foxo1* | For 5'-AAGAGCGTGCCCTACTTCAA-3' |
| | Rev 5'-CTCCCTCTGGATTGAGCATC-3' |
| *Hprt1* | For 5'-TCCTCCTCAGACCGCTTTT-3' |
| | Rev 5'-CCTGGTTCATCATCGCTAATC-3' |
| *Sdha1* | For 5'-CATGCCAGGGAAGATTACAAA-3' |
| | Rev 5'-GTTCCCCAAACGGCTTCT-3' |
| *Pparγ* | For 5'-GCTGTGGGGATGTCTCACAA-3' |
| | Rev 5'-CAGACTCTGGGTTCAGCTGG-3' |
| *Pparα* | For 5'-TTTCCCTGTTTGTGGCTGCT-3' |
| | Rev 5'-TCTGGATGGTTGCTCTGCAG-3' |
| *Pgc-1α* | For 5'-GGACGGAAGCAATTTTTCAA-3' |
| | Rev 5'-TTACCTGCGCAAGCTTCTCT-3' |
| *Nrf1* | For 5'-GATGCTAATGGCCTGGTCCA-3' |
| | Rev 5'-GCTGTCCGATATCCTGGTGG-3' |
| *Nrf2* | For 5'-GAGCAGGACATGGAGCAAGT-3' |
| | Rev 5'-TCTGTCAGTGTGGCTTCTGG-3' |
| *Tfam* | For 5'-TCTGTCTCCTGAGGAAAAGCAG-3' |
| | Rev 5'-ACTTCGTCCAACTTCAGCCA-3' |
| *Mfn1* | For 5'-CTATGCAACCGAGAAGCTGC-3' |
| | Rev 5'-GTTGGCACAGTCGAGCAAAA-3' |
| *Mfn2* | For 5'-ACGCCAGTGAGAAGCTACAG-3' |
| | Rev 5'-GGTGATGTCAACTTGCTGGC-3' |
| *Opa1* | For 5'-GGCAGCAGCTTACAAACACT-3' |
| | Rev 5'-GCTGAACTCGTTTGCCAGTG-3' |
| *Drp1* | For 5'-GAGTGAACTGGTAGGGCAGC-3' |
| | Rev 5'-GACTGGCTCCTTGTAATGCCT-3' |
| *Fis1* | For 5'-ATATGCCTGGTGCCTGGTTC-3' |
| | Rev 5'-AGTCCCGCTGTTCCTCTTTG-3' |
| *Gpx1* | For 5'-TGAACGATCTGCAGAAGCGT-3' |
| | Rev 5'-CAGGTCGGACGTACTTGAGG-3' |
| *Sod2* | For 5'-GTGGGAGTCCAAGGTTCAGG-3' |
| | Rev 5'-ATCCCCAGCAGCGGAATAAG-3' |
| *Prdx3* | For 5'-ACTCCGTTTGGTAAAGGCGT-3' |
| | Rev 5'-AGCTGTTGGACTTGGCTTGA-3' |
| *Pink1* | For 5'-TCCTTGTGGAGTGGGACTCA-3' |
| | Rev 5'-CACCACGCTCTACACTGGAG-3' |
| *D-loop* | For 5'-GACCAACATAACTGTGGTGTCA-3' |
| | Rev 5'-ATTCTTCTCCGTAGGTGCGTCTAG-3' |
| *16srRNA* | For 5'-CCGCAAGGGAAAGATGAAAGAC-3' |

| | |
|---|---|
| | Rev 5'-TCGTTTGGTTTCGGGGTTTC-3' |
| *Nd1* | For 5'-CTAGCAGAAACAAACCGGGC-3' |
| | Rev 5'-CCGGCTGCGTATTCTACGTT-3' |
| *B2m* | For 5'-TGTCAGATATGTCCTTCAGCAAGG-3' |
| | Rev 5'-TGCTTAACTCTGCAGGCGTATG-3' |
| *Hk2* | For 5'-GCCAGCCTCTCCTGATTTTAGTGT-3' |
| | Rev 5'-GGGAACACAAAAGACCTCTTCTGG-3' |
| *Nd4* | For 5'-TAATCGCACATGGCCTCACA-3' |
| | Rev 5'-TTTGAAGTCCTCGGGCCATG-3' |

#### Supplementary Table 2: Antibody list

| Primary Antibody | Supplier | Application & dilution | Catalog # | RRID |
|---|---|---|---|---|
| Anti-FOXP1 antibody (rabbit) | Abcam | WB 1:1000; Co-IP 1:100 | ab16645 | AB_732428 |
| Anti-PGC-1 Alpha antibody | Abcam | WB 1:1000 | ab54481 | AB_881987 |
| Anti-Mitofusin 1 antibody | Abcam | WB 1:1000 | ab104274 | AB_10712138 |
| Anti-Nrf1 antibody | Abcam | WB 1:1000 | ab34682 | AB_2236220 |
| Anti-TFAM antibody | Abcam | WB 1:1000 | ab131607 | AB_11154693 |
| Anti-TTC11/FIS1 antibody | Abcam | WB 1:1000 | ab71498 | AB_1271360 |
| Anti-GAPDH rabbit antibody | Abcam | WB 1:5000 | ab9485 | AB_307275 |
| Anti-GAPDH mouse antibody | Abcam | WB 1:5000 | ab8245 | AB_2107448 |
| COX IV Antibody | Cell Signaling Technology | WB 1:1000 | 4844S | AB_2085427 |
| Foxo1 Rabbit mAb | Cell Signaling Technology | WB 1:1000; Co-IP 1:100 | 2880S | AB_2106495 |
| LC3A/B (D3U4C) Rabbit mAb | Cell Signaling Technology | WB 1:1000 | 12741S | AB_2617131 |
| Mouse IgG1 kappa Isotype control | ThermoFisher Scientific | Co-IP 1:100 | 14-4714-82 | AB_470111 |
| Purified Mouse Anti-OPA1 | BD Biosciences | WB 1:1000 | 612606 | AB_399888 |
| Purified Mouse Anti-DLP1 | BD Biosciences | WB 1:1000 | 611112 | AB_398423 |
| Phospho-DRP1 (Ser637) Antibody | Cell Signaling Technology | WB 1:1000 | 4867S | AB_10622027 |
| Parkin (Prk8) Mouse mAb | Cell Signaling Technology | WB 1:1000 | 4211S | AB_2159920 |
| Purified Mouse Anti-Cytochrome C | BD Biosciences | WB 1:1000 | 556433 | AB_396417 |

| Secondary Antibody | Supplier | Application & dilution | Catalog # | RRID |
|---|---|---|---|---|
| IRDye 800CW Donkey anti- Rabbit | LI-COR Biosciences | WB 1:10000 | 926-32213 | AB_621848 |
| IRDye 680CW Donkey anti- Mouse | LI-COR Biosciences | WB 1:10000 | 926-68072 | AB_10953628 |

#### Supplementary Table 3: CatWalk gait analysis in WT (n = 11) and *Foxp1*$^{+/-}$ mice (n = 6)

| Parameters | WT | *Foxp1*$^{+/-}$ | *p* value |
|---|---|---|---|
| **Basic Locomotion** | | | |
| Run average speed (cm/s) | 24.83 ± 1.74 | 16.27 ± 1.23 | 0.028 (*) |
| Run duration (s) | 1.97 ± 0.15 | 2.99 ± 0.20 | 0.014 (*) |
| Total number of steps | 20.09 ± 0.57 | 25.78 ± 0.88 | 0.001(***) |
| Step sequence number of footfall patterns[a] | 3.97 ± 0.15 | 5.11 ± 0.19 | 0.002 (**) |
| **Step Characteristics** | | | |
| Stride length (cm)[b] | 6.46 ± 0.18 | 5.05 ± 0.17 | 0.001(***) |
| Body speed (cm/s)[c] | 25.25 ± 1.64 | 17.12 ± 1.21 | 0.025(*) |
| Swing speed (cm/s)[d] | 55.52 ± 2.62 | 45.52 ± 2.26 | 0.110 |
| Stand (s)[e] | 0.15 ± 0.01 | 0.19 ± 0.01 | 0.122 |
| Duty cycle (%)[f] | 51.96 ± 1.28 | 58.40 ± 1.11 | 0.011(*) |
| Initial dual stance (s)[g] | 0.02 ± 0.003 | 0.04 ± 0.005 | 0.011(*) |
| Terminal dual stance (s)[g] | 0.02 ± 0.004 | 0.04 ± 0.005 | 0.017(*) |
| Max contact max intensity[h] | 141.65 ± 4.38 | 123.34 ± 3.63 | 0.073 |
| Max contact mean intensity[h] | 65.2 ± 0.91 | 61.30 ± 0.98 | 0.079 |
| Max intensity[i] | 147.53 ± 4.67 | 130.58 ± 4.12 | 0.116 |
| Mean intensity[i] | 68.37 ± 1.07 | 64.76 ± 1.07 | 0.190 |
| Min intensity[i] | 30.28 ± 0.17 | 30.21 ± 0.14 | 0.78 |
| Mean intensity of the 15 most intense pixels[j] | 110.04 ± 3.78 | 97.88 ± 3.16 | 0.190 |
| Print area[k] | 0.23 ± 0.013 | 0.21 ± 0.009 | 0.648 |
| Print length[k] | 0.74 ± 0.013 | 0.71 ± 0.016 | 0.354 |
| Print width[k] | 0.63 ± 0.023 | 0.61 ± 0.023 | 0.813 |
| Support single (%)[l] | 7.29 ± 1.36 | 2.39 ± 0.97 | 0.024 (*) |
| Support diagonal (%)[l] | 72.76 ± 1.10 | 63.30 ± 2.42 | 0.014 (*) |
| Support three (%)[l] | 12.16 ± 1.78 | 24.08 ± 2.75 | 0.011 (*) |
| Support four (%)[l] | 2.86 ± 1.05 | 8.30 ± 1.96 | 0.069 |
| **Gait Variability** | | | |
| Maximum variation (%) | 31.67 ± 3.79 | 60.96 ± 6.55 | 0.005 (**) |
| Body speed variation (%)[m] | 14.43 ± 2.30 | 26.94 ± 3.16 | 0.017(*) |
| **Interpaw Coordination** | | | |
| Phase dispersions[n] | | | |
| RF->LH | 66.99 ± 6.30 | 5.75 ± 0.98 | 0.000(***) |
| LF->RH | 60.37 ± 9.08 | 4.42 ± 1.21 | 0.003(**) |
| RF->RH | 45.89 ± 1.42 | 53.58 ± 0.51 | 0.007(**) |
| LF->LH | 47.63 ± 0.91 | 55.11 ± 0.97 | 0.001(***) |
| LF->RF | 50.23 ± 0.85 | 50.58 ± 0.85 | 0.916 |
| LH->RH | 46.92 ± 1.96 | 49.26 ± 1.17 | 0.553 |
| Couplings[n] | | | |
| RF->LH | 72.31 ± 7.44 | 5.24 ± 0.75 | 0.000(***) |
| LF->RH | 61.11 ± 10.25 | 25.62 ± 10.24 | 0.057 |
| LH->RF | 33 ± 7.68 | 95.53 ± 0.20 | 0.001(***) |
| RH->LF | 30.18 ± 8.79 | 83.66 ± 6.84 | 0.007(**) |
| RF->RH | 46.17 ± 1.24 | 53.99 ± 0.73 | 0.004(**) |
| LF->LH | 46.41 ± 1.51 | 55.66 ± 1.21 | 0.003(**) |
| RH->RF | 54.39 ± 1.06 | 46.06 ± 1.39 | 0.002(**) |
| LH->LF | 53.64 ± 1.52 | 44.59 ± 0.94 | 0.009(**) |
| LF->RF | 49.86 ± 0.92 | 50.47 ± 0.74 | 0.960 |

| | | | |
|---|---|---|---|
| LH->RH | 47.94 ± 1.53 | 48.23 ± 0.42 | 0.887 |
| RF->LF | 49.48 ± 0.75 | 49.64 ± 1.02 | 0.461 |
| RH->LH | 50.88 ± 1.47 | 48.48 ± 1.84 | 0.418 |
| **a.** The step sequence lists the order in which the paws were placed on the glass plate | | | |
| **b.** Distance between successive placements of the same paw | | | |
| **c.** Distance that the animal's body traveled from one initial contact of a specific paw to next divided by the time to travel that distance | | | |
| **d.** Speed of the paw during swing (duration of no contact of a paw with the glass plate) | | | |
| **e.** Duration of contact of a paw with the glass plate | | | |
| **f.** Duty cycle expressed stand as a percentage of step cycle (time between two consecutive initial contacts of the same paw) | | | |
| **g.** Dual stance is the duration of ground contact for both hind paws simultaneously. Initial dual stance is the first time and terminal dual stance is the second time in a step cycle of a hind paw that the contralateral hind paw also makes contact with the glass plate. | | | |
| **h.** Maximum/mean intensity at max contact (the time since the start of the run that a paw makes maximum contact with the glass plate) | | | |
| **i.** Maximum/mean/minimum intensity of the complete paw | | | |
| **j.** Mean intensity of the 15 pixels of a paw with the highest intensity | | | |
| **k.** Print length is the length (horizontal direction) of the complete print. Print width is the width (vertical direction) of the complete paw print. Print area is the surface area (in the Distance Unit you selected in the Preferences) of the complete print. The Print area is by definition at least as large as Max Contact Area. | | | |
| **l.** Standing on single paw, diagonal pair of paws, three or four paws. This displays the relative duration of simultaneous contact with the glass plate of all combinations of paws | | | |
| **m.** Calculated by dividing the absolute difference between the body speed and the average speed of a run | | | |
| **n.** Temporal relationship between placement of two paws within a step cycle: diagonal pairs (RF-LH, LF-RH), ipsilateral pairs (RF-RH, LF-LH) and girdle pairs (LF-RF, LH-RH); the range of circular statistics is between 0 and 100. | | | |

(*$p \leq 0.05$, **$p \leq 0.01$, ***$p \leq 0.001$; three-way ANOVA)

### Supplementary Refences


1. Pitzer C, Kuner R, Tappe-Theodor A. Voluntary and evoked behavioral correlates in neuropathic pain states under different housing conditions. *Mol Pain*. 2016;12:1744806916656635.
2. Castro B, Kuang S. Evaluation of muscle performance in mice by treadmill exhaustion test and whole-limb grip strength assay. *Bio Protoc*. 2017;7:e2237.
3. Hering T, Birth N, Taanman JW, Orth M. Selective striatal mtDNA depletion in end-stage Huntington's disease R6/2 mice. *Exp Neurol*. 2015;266:22-9.
4. He L, Chinnery PF, Durham SE, Blakely EL, Wardell TM, Borthwick GM, et al. Detection and quantification of mitochondrial DNA deletions in individual cells by real-time PCR. *Nucleic Acids Res*. 2002;30:e68.
5. Berkel S, Tang W, Treviño M, Vogt M, Obenhaus HA, Gass P, et al. Inherited and de novo *SHANK2* variants associated with autism spectrum disorder impair neuronal morphogenesis and physiology. *Hum Mol Genet*. 2012;21:344-57.
6. Burstenbinder K, Rzewuski G, Wirtz M, Hell R, Sauter M. The role of methionine recycling for ethylene synthesis in Arabidopsis. *Plant J*. 2007;49:238-49.
7. Chenouard N, Buisson J, Bloch I, Bastin P, Olivo-Marin J-C. Curvelet analysis of kymograph for tracking bi-directional particles in fluorescence microscopy images. 2010 IEEE International Conference on Image Processing2010. p. 3657-60.
8. de Chaumont F, Dallongeville S, Chenouard N, Herve N, Pop S, Provoost T, et al. Icy: an open bioimage informatics platform for extended reproducible research. *Nat Methods*. 2012;9:690-6.
9. Berg S, Kutra D, Kroeger T, Straehle CN, Kausler BX, Haubold C, et al. ilastik: interactive machine learning for (bio)image analysis. *Nat Methods*. 2019;16:1226-32.
